# BMAL1 and YAP cooperate to hijack enhancers and promote inflammation in the aged epidermis

**DOI:** 10.1101/2025.04.22.649967

**Authors:** Júlia Bonjoch, Paloma Solá, Oscar Reina, Thomas Mortimer, Yekaterina A. Miroshnikova, Sara A. Wickström, Camille Stephan-Otto Attolini, Laura Álvarez, Salvador Aznar Benitah, Guiomar Solanas

## Abstract

Ageing is characterised by persistent low-grade inflammation that is linked to impaired tissue homeostasis and functionality. However, the molecular mechanisms driving age-associated inflammation remain poorly understood. The mammalian skin is a clinically relevant target of age-driven inflammation associated with compromised barrier function, inefficient wound healing, elevated oxidative stress, and DNA damage accumulation. Here, we show that upon ageing a previously uncharacterised BMAL1-YAP transcriptional complex is hijacked from chromatin regions associated with homeostatic genes in adult epidermis and redirected to inflammation-related enhancers, amplifying the transcription of their target genes. Independently of its known role as a core circadian clock component, BMAL1 partners with the mechanosensitive transcriptional cofactor YAP at enhancer regions to regulate epidermal identity genes. In contrast, in aged skin, BMAL1-YAP complexes bind to enhancers of inflammation-related genes, co-regulated by NF-KB. Interestingly, aged pro-inflammatory signals from the IL-17 pathway activate YAP in a Hippo-independent manner. These findings unveil a transcriptional mechanism underlying epidermal ageing, linking chromatin dynamics to inflammatory transcriptional programs through BMAL1-YAP-bound enhancer rewiring. By elucidating how ageing reprograms transcriptional networks, our work highlights potential strategies to counteract chronic inflammation and restore tissue homeostasis across age-related loss of functionality.

## Introduction

Organismal regenerative capacity declines with age due to the inability to successfully respond to and resolve stress. This is at least in part driven by chronic, low-grade tissue inflammation^1,2^. During homeostasis, inflammation plays a crucial role in the immune response, helping to defend against pathogens and regulate tissue repair. However, persistent low-grade inflammation during ageing hinders tissue responses to damage due to decreased stem cell function^2–5^. In particular, in mammalian skin, inflammatory responses involve the activation of various immune cells, which contribute to pathogen clearance and tissue repair. Of note, the resolution of this inflammation is essential for effective wound healing and barrier function^6^. Ageing-associated elevated expression of inflammatory mediators impairs stem cell (SC) function and compromises the critical skin barrier function, hair regeneration, and wound healing responses^7–11^.

Previous analyses of aged chromatin architecture indicate global changes in DNA methylation patterns and chromatin mark signatures that eventually lead to chromatin remodelling ^12–15^. Nevertheless, the mechanism by which these changes tune transcriptional programs to impact aged tissue physiology remains poorly understood. Additionally, a new daily rhythmic transcriptional programme is activated in aged interfollicular epidermal (IFE) stem cells, which imposes daily rhythms upon inflammatory mediators^16^. This rewiring of circadian biology is conserved across tissues and organisms and is accompanied by an epigenetic landscape reorganisation^7,16–25^. The circadian core clock machinery is a self-sustained transcription-translation feedback loop that drives organismal rhythmicity. Brain and muscle Arnt-like protein-1 (BMAL1) is one of the main transcription factors of the core clock machinery, driving rhythmic transcription across almost all organs and tissues^22^. In the skin, BMAL1 plays an essential role in maintaining epidermal homeostasis and supporting epidermal progenitors and its loss leads to premature ageing^26–29^. Beyond its circadian clock regulatory role, BMAL1 has been implicated in orchestrating transcriptional responses to stress, suggesting a role in age-associated transcriptional changes^30,31^. However, many BMAL1 targets are transcriptionally arrhythmic, suggesting that it may perform a clock-independent function as a conventional, non-rhythmical transcription factor (TF)^32^.

Mechanotransduction and the circadian clock exhibit a bidirectional interaction. While the circadian core clock machinery interacts with the Hippo pathway^33^, mechanical signals influence circadian clock oscillations in a cell-type-specific manner^34,35^. However, the precise mechanisms underlying the interplay between these two pathways remain unexplored. YAP/TAZ are the main nuclear effectors of the Hippo mechanotransduction pathway. Upon changes in the mechanical environment of a cell, the Hippo pathway becomes inactivated and regulates YAP/TAZ activity by phosphorylation (in the case of YAP, the residue Serine 127). This YAP phosphorylation releases the cytoplasmic retention, eluding subsequent degradation, thereby allowing its transcriptional coactivator function in the nucleus^36^. YAP/TAZ are signalling integrators that reprogram chromatin states and boost enhancer activity^37,38^. In the skin, this canonical function of YAP is essential for its epidermal development and homeostasis during adult life^39,40^. Ageing alters the mechanical properties of skin, including changes in the basement membrane (BM), dermal density and extracellular matrix (ECM) composition^41–44^. Both structures serve as critical niche structures for epidermal SCs, indicating that these structural changes might be affecting SC fitness. Actually, in aged hair follicle stem cells (HFSCs), increased BM stiffness leads to nuclear deformation that triggers transcriptional repression and chromatin remodelling at key stem cell function and activation genes that compromise their function^15^.

Here, we describe a transcription factor (TF) network in control of homeostatic and identity genes operating in mouse skin IFE. We have identified that the core clock component BMAL1 forms a complex with the mechanotransducer Yes-associated protein (YAP), which binds to enhancer elements of genes essential for IFE homeostatic functions. Intriguingly, this BMAL1 enhancer binding is not exclusive to rhythmically expressed genes, highlighting a circadian clock-independent role for BMAL1. Moreover, this regulatory transcription network is hijacked during ageing. In the aged epidermis, the BMAL1-YAP complex binds to enhancer elements of inflammation-related genes, co-regulated by p65, a core transcription factor of the NF-KB pathway. Additionally, we found that YAP signalling is amplified *in vivo* in aged IFE. Mechanistically, we know that YAP is driven not only by mechanical changes (i.e. canonical Hippo activation) but also augmented by tissue inflammatory response. Reducing systemic age-associated inflammation by blocking excessive IL-17 signalling in aged mice^8^ reduces non-canonical YAP activation, suggesting that inflammation lies at the core of age-associated YAP activation.

Our results uncover a new age-related transcriptional axis involving a newly described BMAL1 and YAP complex, which regulates genes defining epidermal identity and function under homeostatic conditions. However, during ageing, this complex is localised to enhancers within the regulatory regions of inflammation-related genes, boosting NF-KB-dependent inflammatory target gene expression. These findings shed light on how age-related transcriptional reprogramming hijacks homeostatic networks, unveiling potential therapeutic targets to mitigate skin ageing and inflammation.

## Results

### Analysis of the Aged Interfollicular Epidermis Diurnal Transcriptome Reveals a Time-of-day Independent Increase in Inflammatory Gene Expression

Generally, transcriptomic analyses are performed at a single time of the day. However, all tissues, including the epidermis, perform specific functions at determined times throughout the day^23^. To avoid biasing our ageing transcriptional analysis by constricting our comparison to one time of the day, as standardly done^45^, we aimed to obtain a more comprehensive view of the differentially expressed genes (DEGs) by being mindful of circadian rhythms. Therefore, we compared rhythmicity pattern-independent expression levels in the aged and adult epidermis over a 24-hour period using the non-rhythmic gene detection in DryR^46^ (Fig. 1a and Supplementary Table 1). Gene ontology (GO) analysis of the upregulated genes in the aged epidermis throughout the day unveiled 35 genes assigned to GO terms related to inflammatory processes (Fig. 1b and Supplementary Table 1). These inflammation-related genes included *Il6st* (coding for gp130), a co-receptor for Interleukin 6 (IL-6), which is involved in acute inflammatory response and wound healing in the epidermis^47,48^ (Fig. 1c and Supplementary Table 1) and genes coding for pro-inflammatory cytokine maturation mediators, such as Caspase 1 and 4 (*Casp1*, *Casp4*)^49^ (Fig. 1c and Supplementary Table 1).

**Figure 1.**
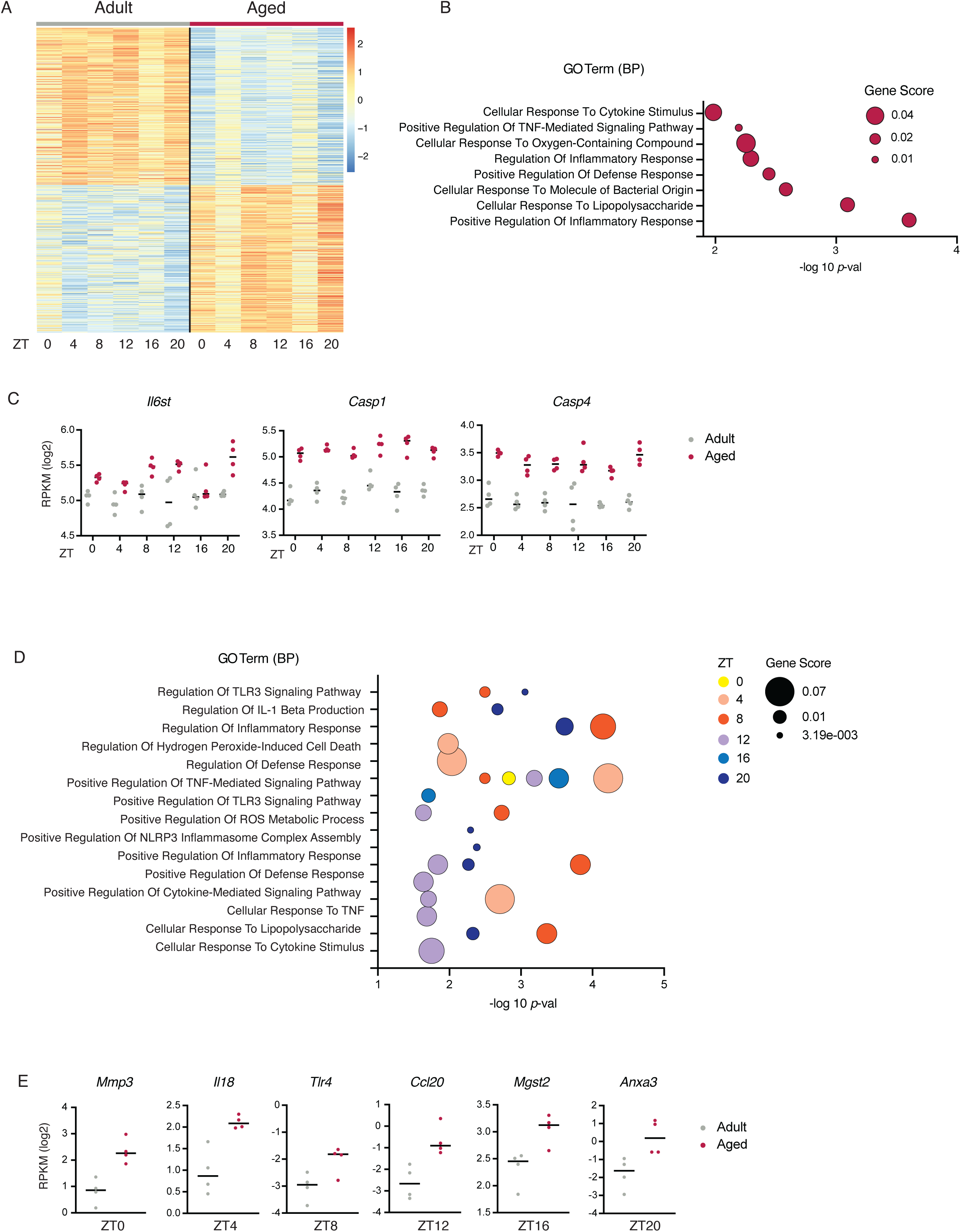
Aged epidermis shows activation of an inflammation-related gene expression program throughout the day. **A.** Heatmap showing expression levels of DEGs in adult and aged epidermis throughout the day (6 ZTs). Transcriptomes of epidermal cells from n=4 adult and n=4 aged mice. **B.** Selected GO BP categories from enrichment analysis of genes up-regulated during ageing irrespective of their rhythmicity pattern. The x-axis represents the -log_10_ of the *p*-value for each depicted category. Size of the dot depicts the gene score (number of genes in the category divided by the total number of genes upregulated). **C**. *Il6st*, *Casp1* and *Casp4* gene expression throughout the day. The y-axis represents the RPKM for each replicate and x-axis shows ZTs. **D.** Selected GO BP categories from enrichment analysis of genes up-regulated during ageing at any given ZT. The x-axis represents the -log_10_ of the *p*-value for each depicted category at different ZTs. Size of the dot depicts the gene score (number of genes in the category divided by the total number of genes upregulated). **E.** *Mmp3*, *Il18*, *Tlr4*, *Ccl20*, *Mgst2* and *Anxa3* gene expression at specific ZTs. The y-axis represents the RPKM for each replicate. GO BP, Gene Ontology Biological Processes; ZT, Zeitgeber time.

Additionally, we performed DEG analysis at every time point extracted -Zeitgeber Time (ZT)-independently to detect genes upregulated at specific times of the day, resulting in 94 genes categorised in GO terms related to inflammatory processes (Extended Data Fig. 1a, Fig. 1d and Supplementary Table 1). These genes include *Mmp3*, a key regulator of cytokine activity by cleavage of their immature form^50^, *Il18*, which plays a role in the chronic inflammatory process of the skin^51,52^, and *Ccl20*, a chemokine-induced by TNF-*⍺*, IL-1 and IL-17A, with a role in psoriasis^53^ (Fig. 1e).

Using both methods, DryR and conventional DEG analysis, a total of 118 genes associated with inflammation functions were detected to be upregulated in the aged epidermis. Notably, of these upregulated genes, only four exhibited rhythmic expression as determined by Meta2D analysis in MetaCycle^54^, confirming that a non-rhythmic pro-inflammatory transcriptional program is upregulated in the aged epidermis (Extended Data Fig. 1a,b and Supplementary Table 1).

### BMAL1 Binds to the Enhancers of Inflammation-Related Genes During Ageing

To investigate BMAL1’s role in age-related inflammation in the epidermis, we assessed its genomic binding in adult and aged epidermis by chromatin immunoprecipitation followed by sequencing (ChIP-seq) during the peak transcriptional activity period of BMAL1, ZT4^32,55^. Importantly, temporal and spatial regulation of gene expression is mediated by cis-regulatory elements (CREs), such as promoters and enhancers, which govern cell identity, homeostasis, and stress responses. These CREs are identified by their enrichment in Histone H3 post-translational modifications, with lysine 4 trimethylation (H3K4me3) marking active promoters, lysine 4 monomethylation (H3K4me1) marking primed enhancers and lysine 27 acetylation (H3K27ac) marking active enhancers^56^. Previous reports in various tissues suggest that BMAL1 preferentially binds to enhancer elements ^57–59^. To gain a deeper understanding of BMAL1 function on the epidermal chromatin, we analysed the abundance of chromatin marks in BMAL1-positive genomic regions and categorised the peaks as associated with promoters, active enhancers, or primed enhancers. Our analysis revealed that while a small fraction of BMAL1 peaks overlapped with active promoter regions, the majority were enriched in H3K4me1-marked enhancer regions, which were further classified as active (H3K27ac-high) or primed/inactive (H3K27ac-low) (Fig. 2a and Extended Data Fig. 2a).

**Figure 2.**
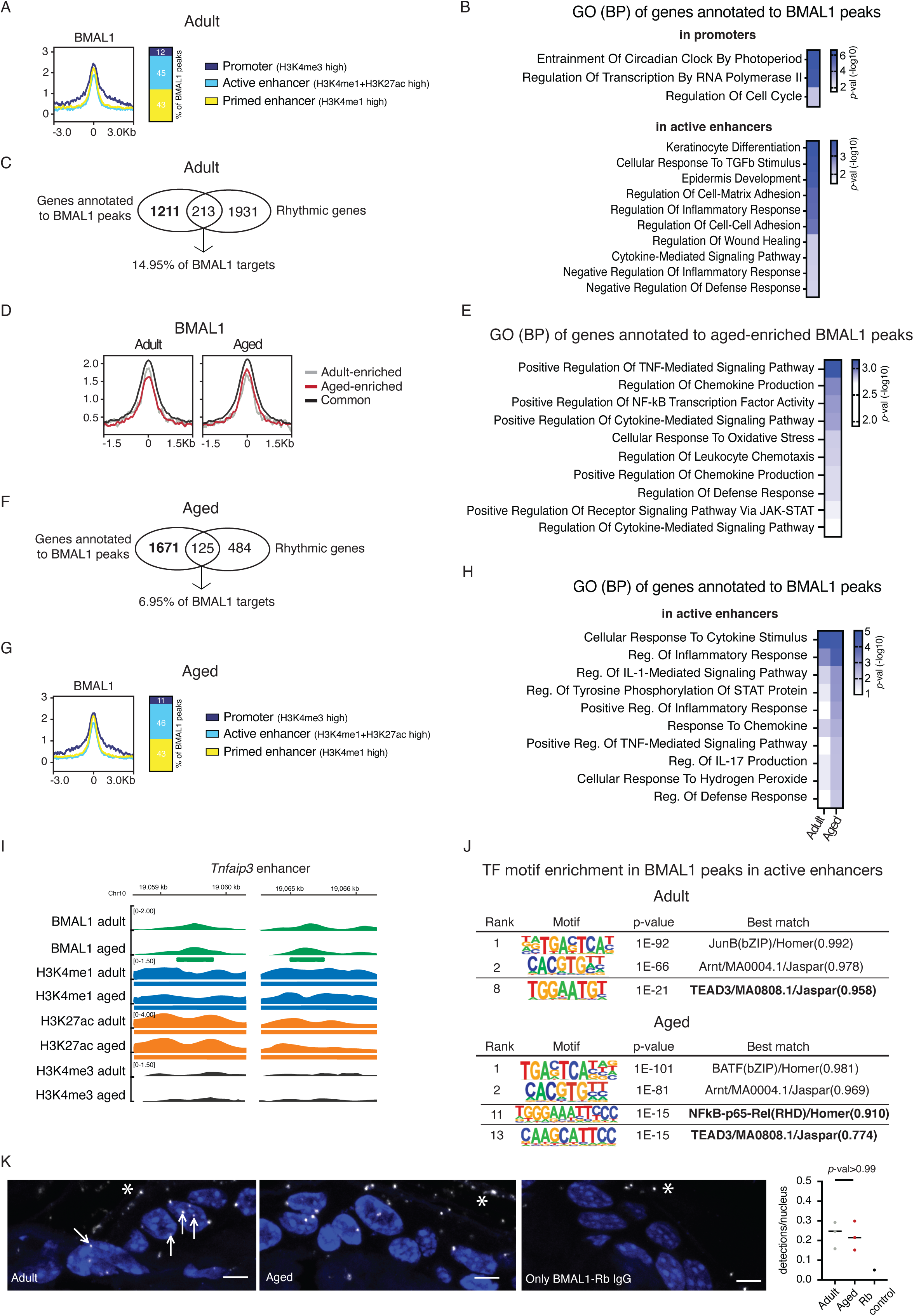
BMAL1 binds to enhancers of inflammation-related genes during epidermal ageing. **A.** BMAL1 ChIP-seq signal ± 3 kb of peak centre at BMAL1 peaks in adult epidermis. K-means (k=3) clustering of BMAL1 peaks was performed based on H3K4me1, H3K27ac, H3K4me3 and BMAL1 signal score. Colours depict BMAL1 signal in each cluster (dark blue=promoter, light blue=active enhancer and yellow=primed enhancer) of BMAL1 peaks. **B.** GO BP enrichment analysis of genes annotated to BMAL1 peaks in promoters and in active enhancers. Only selected GO BP categories are shown. **C.** Comparison of the genes annotated to adult BMAL1 peaks with the rhythmic genes in adult epidermis, obtained by Meta2D with an FDR<0.1. **D.** BMAL1 ChIP-seq signal ± 1.5 kb of peak centre in adult and aged epidermis. K-means (k=3) clustering of BMAL1 peaks was performed based on BMAL1 signal score. Colours depict BMAL1 signal in each cluster (grey=adult enriched, red=aged enriched and black=common) of BMAL1 peaks. **E.** GO BP enrichment analysis of genes annotated to BMAL1 aged-enriched. Only selected GO BP categories are shown. **F.** Comparison of the genes annotated to aged BMAL1 peaks with the rhythmic genes in aged epidermis, obtained by Meta2D with an FDR<0.1. **G.** BMAL1 ChIP-seq signal ± 3 kb of peak centre of BMAL1 peaks in aged epidermis. K-means (k=3) clustering of BMAL1 peaks was performed based on H3K4me1, H3K27ac, H3K4me3 and BMAL1 signal score. Colours depict BMAL1 signal in each cluster (dark blue=promoter, light blue=active enhancer and yellow=primed enhancer) of BMAL1 peaks. **H.** GO BP enrichment analysis of genes annotated to adult and aged BMAL1 peaks in active enhancers. Only selected GO BP categories are shown. **I.** Genomic loci depicting BMAL1 binding to an enhancer region of *Tnfaip3*, one of the inflammation-related genes whose expression is up-regulated in aged epidermis. The thick bars under genomic tracks represent those that have been called a peak for the TF or chromatin mark. Chromosome kilobases are displayed above for identification of the loci. **J.** TF motifs enriched in BMAL1 peaks in active enhancers in adult and aged epidermis, based on *de novo* motif discovery performed with HOMER software. **K.** Representative confocal microscopy images of adult and aged epidermis showing proximity ligation reaction sites (arrows) of BMAL1–YAP interaction in the nucleus. Representative confocal microscopy image of aged epidermis showing a single antibody control for PLA reaction using only BMAL1-Rabbit IgG is also presented. Stars show cornified layer unspecific signal. Scale bars, 5 μm. For enhanced visualization, brightness and contrast were adjusted on these images for visualization purposes exclusively, and equally in both conditions; n=3 adult, n=3 aged and n=1 only BMAL1-Rabbit IgG, mice. On the right, quantification of the number of detections per nucleus. The colour intensity of all heatmaps presented in the figure represents the -log_10_ of the *p*-value for each depicted category. GO BP, Gene Ontology Biological Processes; FDR, False Discovery Rate; TF, transcription factor; PLA, proximity ligation assay.

Annotation of BMAL1 peaks to the nearest genes highlighted functional differences: promoter-bound genes were associated with core circadian functions, whereas enhancer-bound genes regulated epidermal identity and homeostasis (Fig. 2b and Supplementary Table 2). Consistent with other tissues^32^, only a minority of BMAL1 epidermal targets (14.95%) exhibited rhythmic expression in epidermal cells, while being expressed at similar levels, further suggesting that BMAL1 also plays a clock-independent role in regulating epidermal identity and function (Fig. 2c).

A comparative analysis of BMAL1 genomic binding between adult and aged mouse epidermis revealed regions bound by BMAL1 with higher statistical significance in aged epidermal samples (that is, determined as peaks only in aged chromatin samples, termed age-enriched) (Fig. 2d and Extended Data Fig. 2b). Annotation of these age-enriched peaks showed an overrepresentation of genes involved in inflammation (Fig. 2e and Supplementary Table 2). Notably, in aged epidermis, less than 7% of BMAL1 targets showed rhythmic transcription, while being transcribed at similar levels, emphasising its role in non-rhythmic transcription (Fig. 2f and Extended Data Fig. 2c). Also, here BMAL1 was found to localise preferentially to enhancer elements, with enrichment in binding near genes associated with inflammation-related processes (Fig. 2g,h and Extended Data Fig. 2d). As an example, BMAL1 gained binding in two of the enhancers annotated to *Tnfaip3*, a limiting factor of the expression of pro-inflammatory genes in the epidermis^60^ (Fig. 2i).

Additionally, one of the intragenic enhancers of *Usp18*, whose action regulates type I interferon signalling^61^, also gains BMAL1 binding during ageing (Extended Data Fig. 2e). This observation is consistent with the engagement of new enhancer hubs as part of the age-associated transcriptional response^7,14^.

In the epidermis, a coordinated TF circuitry regulates cell identity, homeostasis, and stress responses by modulating chromatin states^62–64^. TF motif enrichment analysis of BMAL1 peaks in active enhancers revealed BMAL1 (*Arntl*) as the second most enriched TF regardless of age. Strikingly, TEAD family factor binding sites were significantly enriched in BMAL1-bound active enhancers in both adult and aged epidermis (Fig. 2j). TEAD TFs mediate YAP/TAZ binding to the DNA, suggesting YAP/TAZ involvement in BMAL1 target gene expression.

To study a possible BMAL1 and YAP functional interaction in epidermal cell chromatin, we performed proximity ligation assay (PLA) on skin sections to identify BMAL1-YAP complex formation ^65^. We observed BMAL1-YAP interactions, depicted by PLA speckles, in both adult and aged IFE (Fig. 2k). In contrast, a single antibody control using the BMAL1-Rabbit IgG alone did not result in the formation of detectable PLA speckle formation inside the nucleus (Fig. 2k). This result suggests that BMAL1 and YAP transcriptional cooperation takes place through complex formation in the nucleus of epidermal cells. We therefore set out to explore whether ageing-related changes to YAP activity in the aged epidermis could underlie the upregulation of an inflammatory gene expression programme.

### YAP Activity Increases in Aged Epidermis

Similarly to the age-associated BM stiffening in the HFSC, we observed elevated BM stiffness in the IFE of aged mice by atomic force microscopy (AFM; Fig. 3a). Consistent with the increased stiffness of aged skin, IFE exhibited higher proportion of epidermal cells positive for the Hippo-dependent active form of YAP (unphosphorylated at serine 127 -unP-S127 YAP)^36^ (Fig. 3b). This finding is consistent with reports of increased YAP activity in other aged tissues, such as luminal epithelial cells in mammary glands and hepatocytes^66,67^. Notably, the increased YAP activation in aged epidermal cells occurred without significant changes in *Yap1* mRNA levels (Fig. 3c).

**Figure 3.**
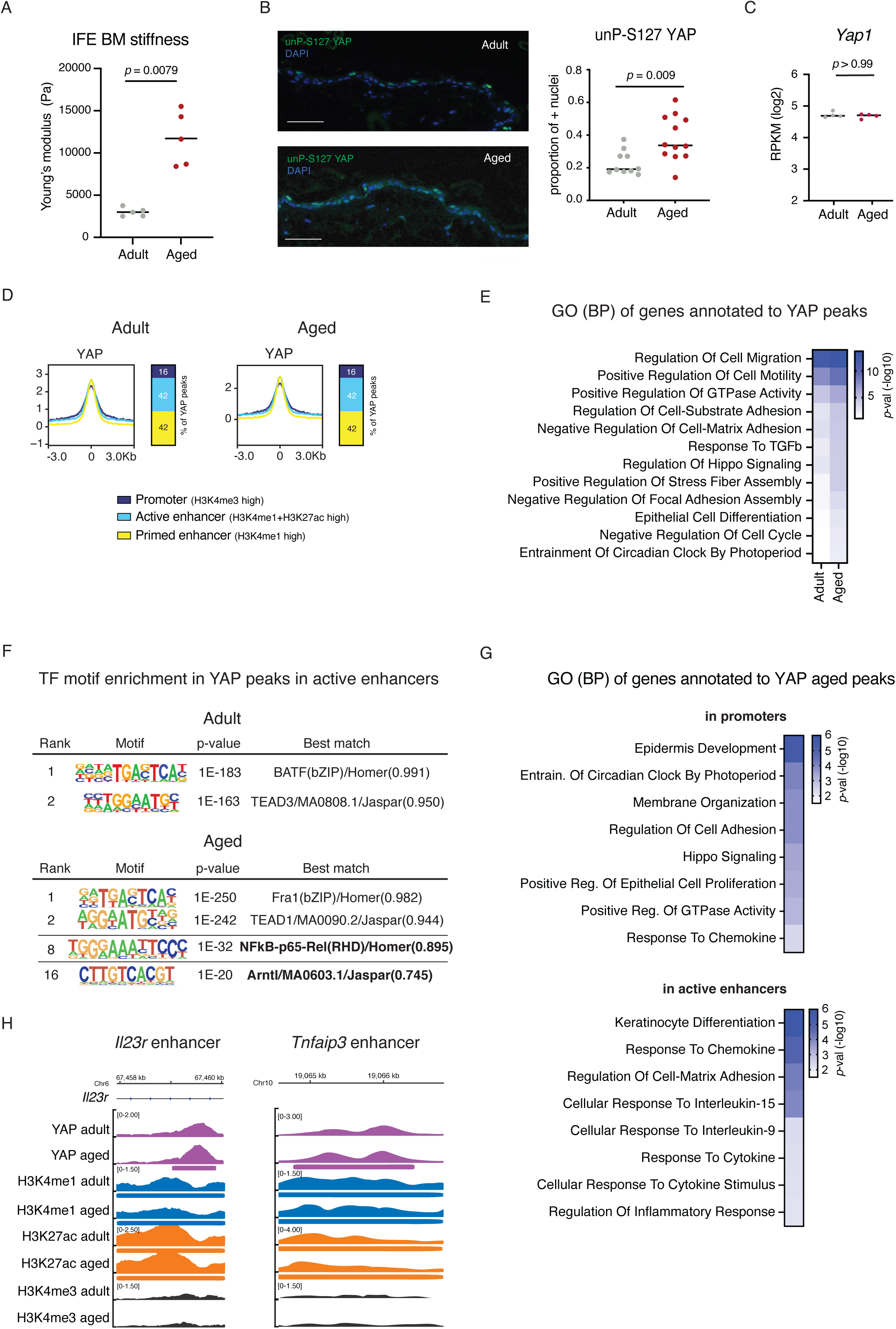
YAP activity is increased in the aged epidermis, impacting gene expression regulation through enhancer binding. **A.** Atomic force microscopy force indentation measurements of IFE BM stiffness. Between 100-140 force curves were obtained for each mouse and means (represented as a dot) were calculated for each mouse. n = 5 mice per age. **B.** Representative images of immunofluorescence staining of unphosphorylated-S127 YAP (unP-S127 YAP) and quantification in adult and aged mouse back skin. Scale bars, 50 μm. For enhanced visualization, brightness and contrast were adjusted on these images for visualization purposes exclusively, and equally in both conditions; n= 10 adult and n= 12 aged mice. Quantification is presented as the proportion of nuclei positive epidermal cells. **C.** *Yap1* gene expression at ZT4. The y-axis represents the RPKM for each replicate. **D.** YAP ChIP-seq signal ± 3 kb of peak centre at YAP peaks in adult and aged epidermis. K-means (k=3) clustering of YAP peaks was performed based on H3K4me1, H3K27ac, H3K4me3 and YAP signal score. Colours depict YAP signal in each cluster (dark blue=promoter, light blue=active enhancer and yellow=primed enhancer) of YAP peaks. **E.** GO BP enrichment analysis of genes annotated to adult and aged YAP peaks. Only selected GO BP categories are shown. **F.** TF motifs enriched in YAP peaks in active enhancers in adult and aged epidermis, based on *de novo* motif discovery performed with HOMER. **G.** GO BP enrichment analysis of genes annotated to YAP peaks in promoters and in active enhancers in aged epidermis. Only selected GO BP categories are shown. **H.** Genomic loci depicting YAP binding to an enhancer region of *Il23r* and an enhancer region of *Tnfaip3*. The thick bars under genomic tracks represent those that have been called a peak for the TF or chromatin mark. Chromosome kilobases are displayed above for identification of the loci. The colour intensity of all heatmaps presented in the figure represents the -log_10_ of the *p*-value for each depicted category. IFE, interfollicular epidermis; BM, basement membrane; ZT, Zeitgeber time; RPKM, Reads Per Kilobase per Million mapped reads; TF, transcription factor.

To understand the transcriptional consequences of increased YAP activity we profiled its binding genome-wide. YAP predominantly binds to enhancer elements, recruiting transcriptional machinery to regulate organ development, size, and stress responses^68,69^. Accordingly, YAP genomic binding in the epidermis showed a preference for enhancer elements, independent of age (Fig. 3d, Extended Data Fig. 3a and Supplementary Table 3). Consistent with the increase in active nuclear YAP, ChIP-seq revealed an increase in the number of YAP peaks in the aged epidermis compared to adult epidermis (Extended Data Fig. 3b and Supplementary Table 3). Notably, YAP aged-enriched targets showed an overrepresentation of mechanotransduction and Hippo signalling pathways by GO analysis (Fig. 3e and Supplementary Table 3), further indicating higher YAP transcriptional activity upon ageing. TF motif enrichment of YAP-bound CREs revealed an overrepresentation of key epidermal lineage regulators, such as AP-1 (Fig. 3f, upper panel). Interestingly, BMAL1 binding sites were significantly enriched exclusively at YAP peaks located in active enhancers in aged epidermis but not in the adult (Fig. 3f, lower panel, *Arntl*), further suggesting a functional relationship between the two TFs in the chromatin of aged epidermal cells.

Moreover, we observed an enrichment of p65-RelA binding boxes in YAP-bound active enhancers only in aged epidermis (Fig. 3f). p65 is one of the transcription factors mediating the NF-KB activity, a key pathway controlling inflammation and immune responses in epidermal cells^70–72^. Consistent, with the age-induced inflammatory phenotype, we observed an enrichment of YAP-bound active enhancers related to inflammatory and cytokine-mediated responses (Fig. 3g). For example, *Il23r* -a mediator of chronic skin inflammatory processes ^73,74^ and an age-specific upregulated gene-gained binding of YAP in a gene-body enhancer in aged epidermal cells (Fig. 3h). Additionally, one of the enhancers annotated to *Tnfaip3*, also gained binding of YAP during ageing (Fig. 3h). Altogether, these data point towards a link between YAP function and inflammation-related processes in the aged epidermis.

### Activation of YAP by Increased Stiffness Is Not Sufficient to Induce Aged-specific YAP Genomic Occupancy

YAP has an established role as a key mediator of enhancer-induced promotion of gene transcription^68,75^. Therefore, we hypothesised that the ECM-induced changes to YAP nuclear translocation and activation in aged skin could create a permissive chromatin state that reprograms the transcriptome of epidermal cells.

To model the ageing-associated BM stiffness increase, we used Collagen XIV (ColXIV) knock-out (KO) mice^76^. Notably, in 1-year-old mice skin, the absence of ColXIV results in a compensatory increase in BM stiffness around HFSCs^15^. AFM measurements in the epidermis indicated an increase in IFE BM stiffness upon depletion of ColXIV (ColXIV KO) already in 47-week-old mice, with stiffness values corresponding to those of aged WT IFE BM (Extended Data Fig. 4a). As predicted, Hippo-dependent YAP activity increased in ColXIV KO epidermal cells already at 47 weeks of age (Extended Data Fig. 4b). However, the absence of ColXIV did not induce major changes in gene expression in mouse back skin epidermal cells as observed by DEG analysis individually at every ZT or irrespective of the time of the day (Extended Data Fig. 4c,d). Combining both methods, a total of 64 genes were found to be upregulated in ColXIV KO epidermal cells when compared to their WT littermates. These DEGs were enriched in GO processes related to barrier formation (i.e. Phosphatidylinositol-3 kinase activity and lipid droplet organisation) (Extended Data Fig. 4e). In contrast, the genes upregulated in aged epidermal cells remained stable upon ColXIV depletion (Extended Data Fig. 4f). This suggests that the increase in BM stiffness driven by disruption of dermal-epidermal junction integrity is not sufficient to induce a transcriptional inflammatory response in epidermal cells.

Epidermal Hippo-dependent YAP activation associated with an increase in BM stiffness in ColXIV KO epidermis is accompanied by a shift in YAP’s genomic location, as analysed by ChIP-seq (Extended Data Fig. 4g and Supplementary Table 4). We compared the genes annotated to peaks called in ColXIV KO and not in WT littermates (stiffness-enriched target genes) to the aged-enriched YAP targets (peaks called exclusively in aged epidermis and not in adult epidermis) to test if stiffness alone was sufficient to relocate YAP to inflammation-related gene’s CREs. Interestingly, stiffness-enriched targets did not recapitulate the presence of YAP in regulatory areas of age-derived stress and inflammation-related genes as shown by GO analysis of the annotated genes and depicted in the example snapshots (Fig. 4a,b).

**Figure 4.**
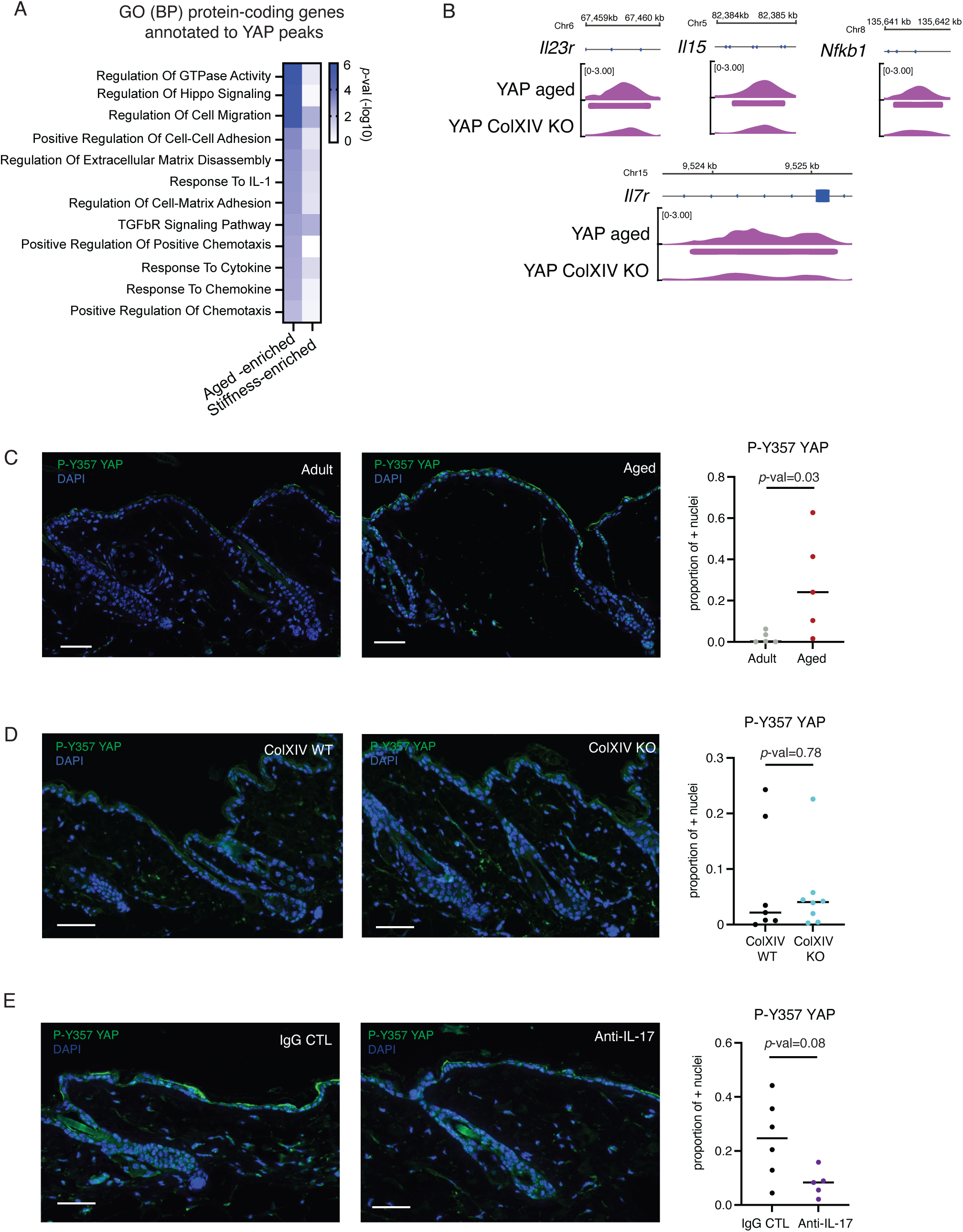
Inflammatory cues promote non-canonical P-Y357 YAP increase in the aged epidermis. **A.** GO BP enrichment analysis of genes annotated to aged-enriched and stiffness-enriched YAP peaks. Only selected GO BP categories are shown. The colour intensity represents the -log_10_ of the *p*-value for each depicted category. **B.** Genomic loci depicting YAP binding to enhancer regions of *Il23r, Il15*, *Il7r* and *Nfkb1*, some of the inflammation-related genes that are bound by YAP in aged epidermis. The thick bars under genomic tracks represent those that have been called a peak for the TF or chromatin mark. Chromosome kilobases are displayed above for identification of the loci. **C.** Representative images of immunofluorescence staining of phosphorylated-Y357 YAP (P-Y357 YAP) and quantification in adult and aged mouse back skin. Scale bars, 50 μm. n= 5 adult and n= 5 aged mice. **D.** Representative images of immunofluorescence staining of P-Y357 YAP and quantification in ColXIV WT and ColXIV KO mouse back skin. Scale bars, 50 μm. n= 7 ColXIV WT and n= 8 ColXIV KO mice. **E.** Representative images of immunofluorescence staining of P-Y357 YAP and quantification in anti-IL-17 and control IgG aged mouse back skin. Scale bars, 50 μm. n= 5 anti-IL-17 and n= 6 control IgG mice. All quantifications are presented as the proportion of epidermal cell nuclei positive for staining. For enhanced visualization, brightness and contrast were adjusted on these images for visualization purposes exclusively, and equally in both conditions that were being compared. GO BP, Gene Ontology Biological Processes

### Non-canonical YAP Activation in the Aged Epidermis

As increased stiffness did not recapitulate YAP genomic occupancy of CREs of age-associated upregulated genes in mouse epidermis, we hypothesised that a second cooperative signalling layer is required to fully activate YAP-mediated transcription and cause its genomic relocation in response to ageing cues.

YAP activity is regulated through post-translational modifications, mainly tyrosine and serine phosphorylation^77^. The Hippo pathway is the canonical regulatory pathway for YAP activity through PTM modulation^36^. Nevertheless, other PTMs regulate YAP activation status in a Hippo-independent non-canonical manner, such as phosphorylation of Tyrosine 357 (P-Y357) by Src family kinases, which leads to YAP transcriptional activation^78–80^. During healing after intestinal mucosal injury, Interleukin 6 (IL-6) signalling induces P-Y357 YAP in response to inflammation through SRC-family kinases (SFK)-mediated phosphorylation, promoting YAP stabilisation and nuclear localisation^78^.

To understand if ageing-associated chronic low-grade inflammation leads to the increase of YAP activity in the aged epidermis by non-canonical PTM changes, we stained skin sections with a P-Y357 YAP-specific antibody. Interestingly, in the aged epidermis, there was a higher proportion of cells with positive P-Y357 YAP nuclei pointing towards an increase in non-canonically activated YAP (Fig. 4c).

Next, we sought to understand the origin of the age-associated signalling underlying the increase in non-canonical YAP activation in the epidermis. As shown above, the increase in BM stiffness alone was not enough to increase P-Y357 YAP in mouse epidermis, as shown by comparable P-Y357 YAP signal in ColXIV KO and WT IFE of 47-week-old mice (Fig. 4d). We therefore assessed if ageing-associated inflammation could be at the root of P-Y357 YAP increase in aged mouse epidermis. IL-17 is a pro-inflammatory cytokine involved in anti-microbial response and the development of autoimmune diseases, including those of the skin, such as psoriasis and alopecia areata^74,81,82^. Aged mouse dermis contains a higher proportion of IL-17-producing lymphoid cells, compared to their adult counterparts^8^. Moreover, *in vivo* anti-IL-17 treatment during ageing reduces inflammation-related gene expression in the epidermis, along with a delay in skin ageing trait acquisition, highlighting IL-17 central role in skin ageing^8^. We hypothesised that ameliorating aged epidermal inflammation by reducing exacerbated IL-17-mediated signalling could reduce YAP activity. While Hippo-dependent YAP activation did not respond to IL-17 blocking treatment (Extended Data Fig. 4h), the non-canonical activating phosphorylation on Y357 decreased significantly upon anti-IL17 treatment in aged mice epidermis (Fig. 4e).

These results suggest that in addition to a Hippo-dependent YAP permissive state due to ECM changes in the aged epidermis, a second layer of activation caused by pro-inflammatory cues leads to ageing-associated YAP activation. We hypothesised that this increase in active YAP nuclear levels might serve as a platform to promote gene expression in collaboration with other TFs.

### BMAL1 and YAP Bind to Enhancers of Inflammation-Related Genes in Aged Epidermis

Considering the predicted overlap of BMAL1 and YAP in active enhancers of upregulated inflammation-related genes in aged epidermis, we decided to further characterise the functional interaction between the two TFs. We observed a statistically significant genomic overlap between BMAL1 and YAP chromatin binding sites regardless of age using a permutation test^83^ (Fig. 5a and Extended Data Fig. 5a,b). These overlapping sites are predominantly located in enhancer regions, with similar proportions between adult and aged samples (Fig. 5b and Extended Data Fig. 5c). Notably, in aged epidermis, genes annotated to these enhancer regions show a greater enrichment for inflammatory processes, suggesting that BMAL1 and YAP could coordinate the transcription of these genes in aged samples by relocating to inflammation-related gene CREs (Fig. 5c and Supplementary Table 5). Some of the BMAL1, YAP and BMAL1-YAP age-enriched peaks were located in the vicinity of CREs of the age-specific inflammation-associated upregulated genes (Fig. 1), such as *Casp1*, *Casp4*, *Il33, Ptgs2, Tnfaip3, Usp18 and Egr1* (Fig. 5d). In some instances, genes that were BMAL1 or YAP targets in adult epidermis, gained BMAL1-YAP peaks at new enhancer regions in the aged IFE cells (Fig. 5e,f). These findings suggest a coordinated interplay between BMAL1 and YAP in enhancing the transcriptional landscape of inflammation-related genes during ageing, potentially through shared occupancy of enhancer regions.

**Figure 5.**
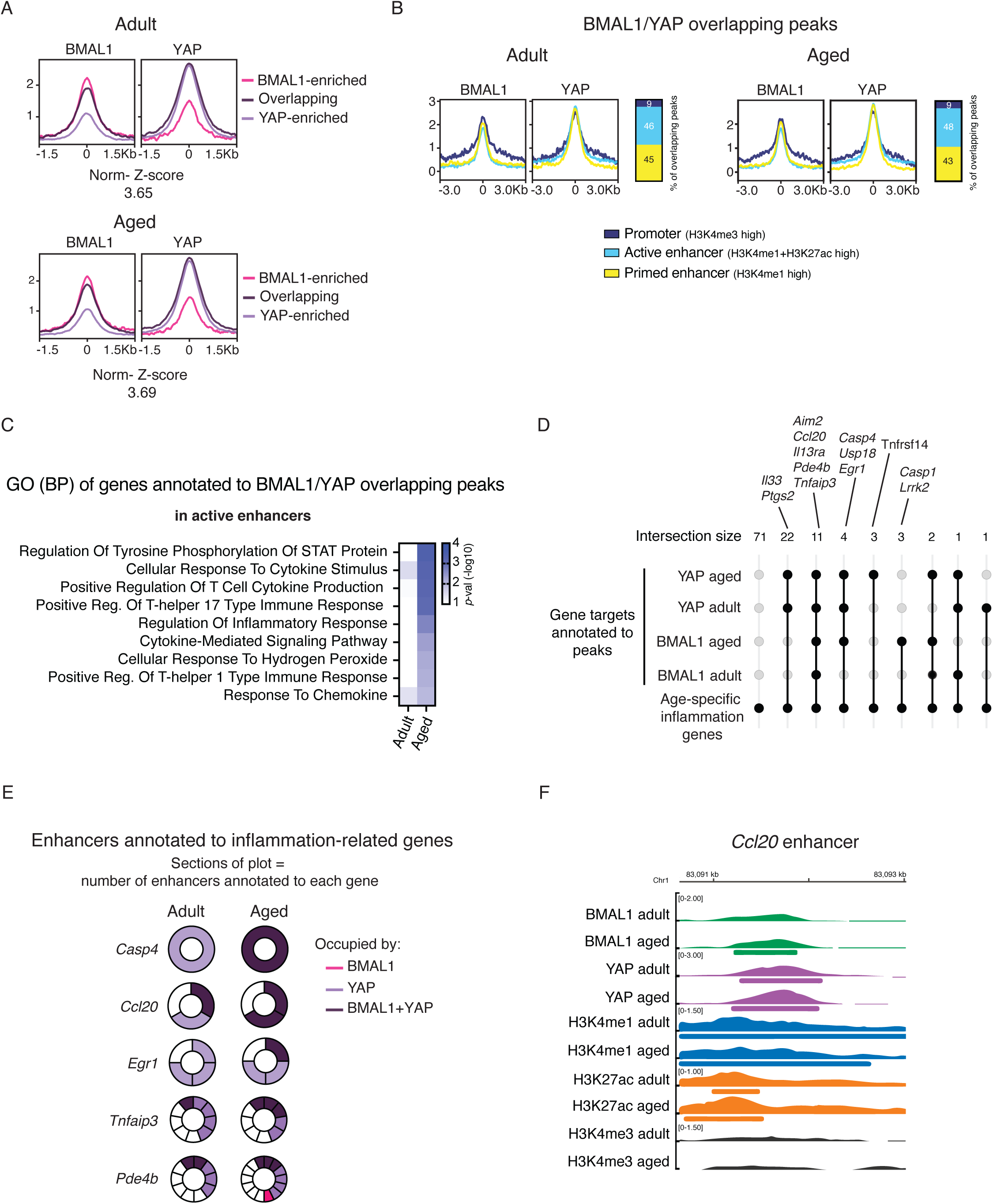
BMAL1 and YAP complex bind to enhancers of key inflammation-related genes whose expression is increased in the aged epidermis. **A.** BMAL1 and YAP ChIP-seq signals ± 1.5 kb of peak centre in adult (upper) and aged (lower) epidermis. K-means (k=3) clustering of BMAL and YAP peaks was performed based on their signal score. Colours depict the signal of the TF in each cluster (pink=BMAL1 enriched, dark purple=overlapping signal, light purple=YAP enriched) of BMAL1 and YAP peaks. **B.** BMAL1 and YAP ChIP-seq signals ± 3 kb of peak centre at BMAL1 and YAP overlapping peaks in adult (left) and aged (right) epidermis. K-means (k=3) clustering of BMAL1 and YAP overlapping peaks was performed based on H3K4me1, H3K27ac, H3K4me3, BMAL1 and YAP signal score. Colours depict the signal of the TFs in each cluster (dark blue=promoter, light blue=active enhancer and yellow=primed enhancer) of BMAL1 and YAP overlapping peaks. **C.** GO BP enrichment analysis of genes annotated to BMAL1 and YAP overlapping peaks in active enhancers in adult and aged epidermis. Only selected GO BP categories are shown. The colour intensity represents the -log_10_ of the *p*-value for each depicted category. **D.** Upset plot representing the number of shared genes between genes annotated to BMAL1 peaks in adult and aged, genes annotated to YAP peaks in adult and in aged, and inflammation genes up-regulated during ageing (5 sets in total). Only intersections with the inflammation gene set are visualised. The above relevant genes of some intersections are highlighted. **E.** Graphical representation of enhancers from chosen inflammation-related genes. Each segment of the circle represents an enhancer annotated to the gene. Colours depict occupancy of an enhancer by a TF (pink=BMAL1, light purple=YAP and dark purple=BMAL1 and YAP). **F.** Genomic loci depicting BMAL1 and YAP binding to an enhancer region of *Ccl20*, one of the inflammation-related genes whose expression is up-regulated in aged epidermis. The thick bars under genomic tracks represent those that have been called a peak for the TF or chromatin mark. Chromosome kilobases are displayed above for identification of the loci. TF, transcription factor; GO BP, Gene Ontology Biological Processes.

### BMAL1–YAP Co-Regulated Enhancers Boost Transcription of p65 Target Genes

We consistently found p65-NF-KB binding sites enriched in aged active enhancers bound by BMAL and YAP (Fig 2j and Fig 3f). We also found that the aged-enriched YAP peaks that were absent in the stiffness-enriched samples (ColXIV KO) showed an overrepresentation of p65-NF-KB binding motifs (Extended Data Fig. 5d). The NF-kB signalling pathway is a molecular cascade that regulates immune response, inflammation, cell survival and stress responses, through NF-KB transcriptional mediators^84^. Among the transcriptionally active units, p65 has been shown to have a key role in the expression of inflammation-related genes during epidermal ageing^8^. Therefore, we further dissected the role of p65 in BMAL1-YAP-bound inflammation-related ageing-associated upregulated genes.

To understand the relevance of p65-mediated transcription in ageing-associated inflammation-related BMAL1-YAP targets genes, we analysed p65 genomic binding in adult and aged epidermal cells by ChIP-seq (Fig. 6a). Interestingly, we identified a higher number of p65-bound targets in aged epidermal cells than in their adult counterparts (Fig. 6a, Extended Data Fig. 6a and Supplementary Table 6). Out of these aged-enriched targets, we observed an overrepresentation of targets involved in NF-KB signalling and inflammatory processes (i.e. *Tlr5*, *Fadd*, *Il22ra2*), as well as ageing-associated stress (i.e. unfolded protein response and oxidative stress), and epidermal homeostatic processes (i.e. cell cycle progression and epidermal identity) (Fig. 6b and Supplementary Table 6). This suggests that p65 could be a transducer of aged-related stress participating in the transcriptional response elicited by aged microenvironmental cues.

**Figure 6.**
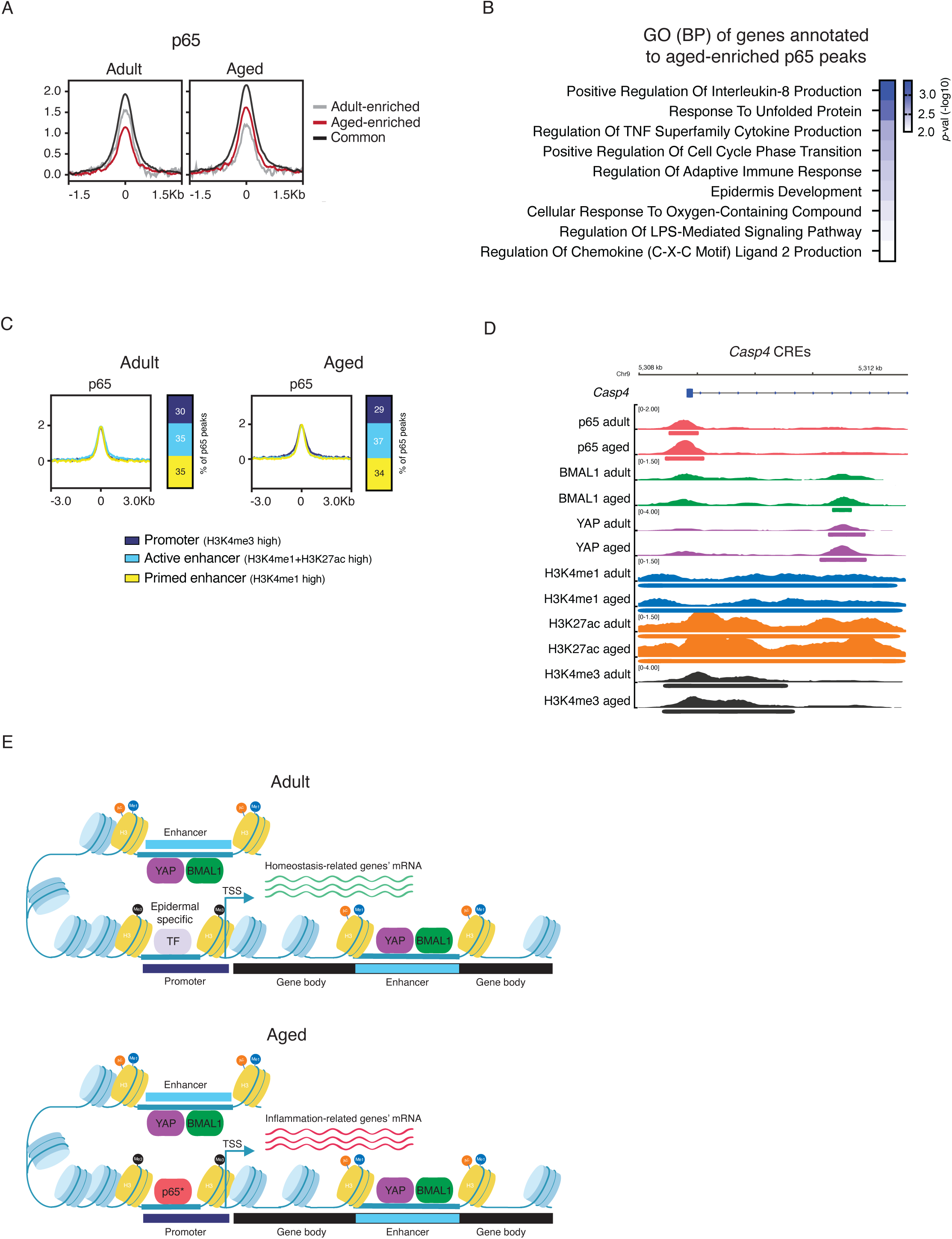
BMAL1 and YAP complex binding to inflammation-related enhancers during ageing supports the expression of p65-target genes. **A.** p65 ChIP-seq signal ± 1.5 kb of peak centre in adult and aged epidermis. K-means (k=3) clustering of p65 peaks was performed based on p65 signal score. Colours depict p65 signal in each cluster (grey=adult enriched, red=aged enriched and black=common) of p65 peaks. **B.** GO BP enrichment analysis of genes annotated to aged-enriched p65 peaks in the epidermis. Only selected GO BP categories are shown. The colour intensity represents the -log_10_ of the *p*-value for each depicted category. **C.** p65 ChIP-seq signal ± 3 kb of peak centre at p65 peaks in adult (left) and aged (right) epidermis. K-means (k=3) clustering of p65 peaks was performed based on H3K4me1, H3K27ac, H3K4me3 and p65 signal score. Colours depict p65 signal in each cluster (dark blue=promoter, light blue=active enhancer and yellow=primed enhancer) of p65 peaks. **D.** Genomic loci depicting p65, BMAL1 and YAP binding to a CRE region of *Casp4*, one of the inflammation-related genes whose expression is up-regulated in aged epidermis. The thick bars under genomic tracks represent those that have been called a peak for the TF or chromatin mark. Chromosome kilobases are displayed above for identification of the loci. **E.** Graphical representation of the proposed epidermal TF network model that rewires gene expression during ageing. In the adult epidermis, BMAL1 and YAP bind to enhancers of homeostatic-related genes, that in cooperation with epidermal-specific TF, maintain identity and homeostasis. In the aged epidermis, these two TFs gain binding to enhancers of inflammation-related genes, including genes whose transcription is p65-NF-kB-dependent. TF, transcription factor; GO BP, Gene Ontology Biological Processes; CRE, Cis-Regulatory Element.

A detailed analysis of p65 genomic feature binding demonstrated that it has a more prominent binding to promoters than BMAL1 or YAP, approximately a 3-fold increase compared to the overlapping BMAL1-YAP peaks (Fig. 6c, Extended Data Fig. 6b, compared to Fig. 2a, 2g, 3e). This preference for promoters did not change with age, remaining stable during epidermal ageing. Notably, the presence of p65 in CREs of age-specific upregulated inflammation-related genes bound by BMAL1-YAP did not overlap with that of BMAL1 and YAP. While p65 had a higher presence in promoters and transcription start sites (TSSs), BMAL1 and YAP binding was more biased towards enhancers, as shown in the snapshot example of *Casp4* CRE (Fig. 6d).

Collectively, these data suggest that in adult epidermis, during homeostatic conditions, BMAL1 and YAP co-ordinately bind to enhancers of identity and SC maintenance genes. During ageing, this TF complex is then directed to inflammation genes, many of which are regulated by p65/NF-KB-binding, increasing their transcription in the aged epidermis (Fig. 6e). Consequently, these changes in gene expression, which are boosted by BMAL1 and YAP action, might contribute to the transcriptional rewiring and loss of function of epidermal cells during ageing.

## Discussion

During ageing, tissues and organs shift their transcriptional programs, transitioning from a homeostatic into a pro-inflammatory state, affecting SC behaviour and tissue regeneration ^2,5,45^. However, most studies have focused on one specific time point, potentially introducing biases towards a particular set of functions. Here, we present a comprehensive transcriptomic analysis covering a range of time points throughout the day. The causes of the age-associated transcriptional switch have been addressed from multiple angles, considering both intrinsic and extrinsic signals^85–88^. Recent studies show that the epigenome changes during ageing, suggesting that epigenetic modulators play a critical role in dictating age-related transcription^7^. Notably, age-associated external stimuli impact chromatin dynamics and, consequently, gene expression by modulating the activity and accessibility to target CREs of key epidermal genes^15,89^. In particular, the aged-derived microenvironmental cues have a great impact on epidermal SC fitness by altering their transcriptomes^8,9,90^. Nevertheless, the molecular players driving this transcriptional switch remain to be fully elucidated.

Our results reveal that a newly identified complex formed by BMAL1 and YAP binds to the CREs of inflammation-related genes to upregulate their expression in the aged mouse epidermis. This finding uncovers an unexpected role for BMAL1 and YAP as ageing-stress transducers, distinct from their canonical roles in the core circadian clock and mechanotransduction pathways, respectively. Age-associated microenvironmental changes in the epidermis cause a shift in BMAL1 and YAP function, repurposing them to elicit a stress response to help epidermal cells adapt to ageing-related challenges. Specifically, we found that BMAL/YAP complexes in aged chromatin are associated with enhancers of genes typically regulated by the NF-KB pathway. This suggests that BMAL1 and YAP act as transcriptional amplifiers of NF-KB signalling in the aged mouse epidermis. Furthermore, studies analysing the transcriptome rewiring of aged human epidermis confirm the upregulation of some of these inflammation-related genes (i.e. *Nfkbia*, *Tnfaip3* and *Il18*)^10^, suggesting that this phenomenon is common across mammalian species. Notably, BMAL1 and YAP were bound not only to pro-inflammatory genes (*Casp1*, *Casp4*, *Il6st*) but also to genes involved in inflammation resolution (*Tnfaip3* and *Usp18*). This dual role suggests a regulatory feedback loop that could prevent excessive NF-KB activation, thereby mitigating potentially irreversible tissue damage.

BMAL1 is the only non-redundant component of the core circadian clock. Its transcriptional targets are tissue-specific, as are the clock output genes, ensuring coordination of daily physiological functions across tissues. Notably, a common feature observed in many tissues is that some BMAL1 target genes are transcribed out of phase with BMAL1 DNA-binding timing, and, strikingly, the majority of target genes are not rhythmically transcribed^32^. Our findings in adult and aged epidermis align with this observation. These results suggest that BMAL1 likely has functions beyond its role as core circadian clock TF^29,91^. The absence of BMAL1 induces premature ageing in many of the tissues analysed in a manner that is hardly attributable to the circadian clock exclusively. Therefore, a detailed study of the circadian clock-independent functions of BMAL1 is needed to understand the relationship between these divergent, and perhaps complementary, functions.

Altered mechanical information of tissue-resident SC niches, including those occurring during ageing, induces defects in tissue progenitor function^92–96^. These studies suggest that a global detrimental response to ageing-associated aberrant mechanical inputs impacts SC fitness. Interestingly, the impact of increased YAP activity during ageing appears to be cell-type specific: in the dermis, lower YAP activity protects fibroblasts from senescence and excessive inflammation^67^, whereas in epithelial tissues and hepatocytes, ageing induces higher YAP activity^66,67^. This differential regulation may reflect the distinct functions of these tissues, with epithelia serving as a physical barrier and an immunologically active interface essential for organismal defence. In the aged epidermis, we show that YAP canonical Hippo-dependent activation alone is insufficient to drive age-specific transcriptional rewiring. Instead, a second layer of non-canonical, inflammation-dependent activation of YAP takes place in the aged epidermis. Previous reports have linked elevated YAP non-canonical activity to pro-inflammatory scenarios, such as cancer and regenerative responses^78,97–99^. Notably, in the intestinal epithelium, gp130 -co-receptor of Interleukin 6 (IL-6) encoded by *Il6st* and upregulated in the aged epidermis-activates Src family kinases, which mediate YAP’s non-canonical activation, promoting mucosal healing following injury^78^. Similarly, we found that ageing-associated inflammatory cues, such as excessive IL-17 signalling, contribute to the non-canonical activation of YAP in the epidermis. IL-17-derived signalling from aged dermis impacts epidermal gene expression mediated by p65/NF-KB^8^. In this sense, we propose that a global decrease in ageing-derived inflammatory cues impacts BMAL1-YAP-p65 mediated transcription by reducing YAP non-canonical activation.

Our results uncover a novel cooperative role for BMAL1 and YAP in the aged epidermis, where they function as stress-responsive transcriptional regulators driving an inflammation-related gene expression program. This shift in function suggests that epidermal cells repurpose existing transcriptional networks to adapt to age-associated stressors. This study expands our understanding of the molecular circuitry underlying age-associated transcriptional rewiring in the epidermis and provides a foundation for future investigations into therapeutic strategies aimed at mitigating age-related tissue dysfunction.

## Methods

### Mouse handling and husbandry

Mice were housed in monitored rooms under 12-hour light/dark cycles and specific-pathogen-free conditions. Animal facility temperature was maintained between 20 and 24 °C and humidity ranged between 45 to 65%. Handling of animals was done following the ethical regulations and guidelines of *Parc Científic de Barcelona* and *Generalitat de Catalunya* governmental authority. All procedures and experimental endpoints were evaluated and approved by the Ethical Committee for Animal Experimentation of *Generalitat de Catalunya* (approval reference no.10712). All mice used were C57BL/6J strain. Wild-type mice were used for the physiological ageing experiments with an age of 80-90 weeks old. Their counterpart adult control mice were between 18 and 25 weeks old. Aged mice were either bred in-house or purchased from Charles River as retired C57BL/6J breeder females, and were kept in the animal facility at *Parc Científic de Barcelona* until they reached the desired age. Adult mice were either bred in-house or purchased from Charles River to have sex-matching cohorts to the aged ones. Both females and males were used for all experiments. Generally, mice were sacrificed 4 hours after lights were turned on (Zeitgeber time 4 -ZT4), unless otherwise stated, to coincide with BMAL1 binding to the chromatin^32^. For daily rhythmic transcriptomic analysis, mice were sacrificed at 4h-apart time points across the dark/light cycle. To avoid interference with the results, mice with signs of skin inflammation (including dermatitis, wounds or redness) were excluded from the experiments.

*Col14a1*^−/-^ transgenic mice (ColXIV KO) were imported from Dr Edgar M Espana’s laboratory at the University of South Florida, and mice were created at Dr Birk’s laboratory at the University of South Florida^76^. ColXIV KO mouse colony was expanded and maintained in the animal facility at *Parc Científic de Barcelona* until they reached the desired age.

### Epidermal cell isolation for single-cell suspension

Mice were sacrificed, shaved and whole-torso skin was obtained. Hypodermal fat was removed with the use of a scalpel followed by two washes in PBS. Then, skins were floated (with the dermal side down) in 0.08% trypsin (Trypsin 1:250, 27250-018, Gibco) in PBS for 30 min for females and 40 min for male mice skins at 37°C. The epidermis was separated from the dermis with a scalpel, and only the epidermis was further processed. The epidermis was kept in a small volume of Calcium-chelated FBS to inactivate trypsin activity. For epidermal cell isolation, the epidermis was mechanically dissociated using a Mcllwain Tissue Chopper (The Mickle Laboratory Engineering Co.). Next, dissociated cells were strained through a 100-µm filter and then through a 40-µm strainer to obtain single-cell suspensions. Finally, the cell suspension was centrifuged at 300 xg at 4°C to pellet down the epidermal cells.

### Bulk RNA sequencing library preparation and sequencing

Cells in single-cell suspensions were frozen at -80°C in 1 ml of Trizol (15596018, Invitrogen) for posterior RNA isolation. RNA was extracted from epidermal cell pellets frozen in Trizol using the NZY Total RNA Isolation Kit (MB13402, NZY Tech). Purified RNA was processed for mRNA-seq with Illumina sequencing technology.

For the adult *vs* aged comparison, libraries were prepared using the TruSeq Stranded Total RNA Library Prep Kit with Ribo-Zero Human/Mouse/Rat Kit (RS-122-2201/2202, Illumina) according to the manufacturer’s protocol, using 150 to 300 ng of total RNA; ribosomal RNA depletion, RNA was then fragmented for 4.5 min at 94°C. The remaining steps were followed according to the manufacturer’s instructions. Final libraries were analysed on an Agilent Technologies 2100 Bioanalyzer system using the Agilent DNA 1000 chip to estimate the quantity and validate the size distribution; libraries were then quantified by qPCR using the KAPA Library Quantification Kit KK4835 (07960204001, Roche) before amplification with Illumina’s cBot. Finally, libraries were sequenced on the Illumina HiSeq 2500 sequencing system using paired-end 50-base pair (bp)-long reads.

For the ColXIV KO vs WT comparison, Libraries were prepared using the TruSeq stranded mRNA Library Prep (20020595, Illumina) according to the manufacturer’s protocol, to convert total RNA into a library of template molecules of known strand origin and suitable for subsequent cluster generation and DNA sequencing. Briefly, 500 ng of total RNA were used for polyA-mRNA selection using Oligo-dT beads, and two rounds of purification were performed. During the second elution of the mRNA, this was fragmented under elevated temperature and primed with random hexamers for cDNA synthesis, which was performed using reverse transcriptase (SuperScript II; 18064-014, Invitrogen). Then, second strand cDNA was synthesised, incorporating dUTP in place of dTTP and generating blunt-ended ds cDNA. A single ‘A’ nucleotide was added to the 3’ ends of the blunt fragments (A-tailing) and immediately afterwards the Truseq adapter was ligated. Finally, PCR selectively enriched those DNA fragments that had adapter molecules on both ends. The PCR was performed using Unique Dual Indexes and the master mix provided with the kit. All purification steps were performed using AgenCourt AMPure XP beads (A63882, Beckman Coulter). Final libraries were analysed using Bioanalyzer DNA 1000 or Fragment Analyzer Standard Sensitivity (5067-1504 or DNF-473, Agilent) to estimate the quantity and validate the size distribution and were then quantified by qPCR using the KAPA Library Quantification Kit KK4835 (07960204001, Roche). Libraries were sequenced 1 X 51+10 +10 bp-long reads on Illumina’s NextSeq2000.

### Differential expression analysis of bulk RNA-seq data

FastQ files with reads were aligned to the mm10 reference genome using STAR aligner version 2.5.2b^100^ with default settings. Expected read counts for Entrez mm10 annotation were obtained with the featureCounts function from the Rsubread package version 1.32.4 with options countMultiMappingReads=TRUE, allowMultiOverlap=TRUE, isPairedEnd=TRUE^101^. Read counts were input for DEG analysis. Differentially expressed genes (DEGs) were obtained with DESeq2 1.22.2^102^. Genes with lfcShrink |FC|>1.5 (type=normal) and Benjamini-Hochberg adjusted *P*-value<0.1 were considered as differentially expressed.

### DryR function

To identify genes that were differentially expressed in the epidermis irrespective of the time of day in adult versus aged mice and wild type versus ColXIV KO mice, we applied the DryR algorithm^46^. Log_2_ transformed RPKM (reads per kilobase per million mapped reads) values were used as input, and genes were pre-filtered to remove those with an expression standard deviation across all time points = 0 or expression in at least one replicate ≤ 0.1 RPKM (unexpressed genes). The function ‘drylm()’ was then applied to run the algorithm. The resulting output was then filtered (mean model = 2, BICW mean model 2’ 0.95, Log_2_ fold change 2’ 0.25, Log2 expression 2’ 1 in at least one condition) to generate a set of differentially expressed genes for downstream analysis.

### Identification of rhythmic genes with MetaCycle meta2D

Rhythmic gene expression was evaluated independently for adult and aged epidermal cells with the MetaCycle tool 1.2.0.^54^. DESeq2 log2 normalised counts at Entrez gene level (featureCounts/RSubread 1.32.4) were used, and CycMethods (ARS, JTK and LS) was performed, with the meta2d function with minper20 and maxper24 options. Method ARS was rejected due to being unable to work with biological and/or technical replicates. Meta2D results are an integration of JTK and LS results. Transcripts with a BH q value less than 0.1 were considered rhythmic.

### Chromatin extraction and chromatin immunoprecipitation

For ChIP followed by sequencing (ChIP-seq) primary epidermal cells in a single-cell suspension were used as starting material. For ChIP of Histone modification marks, 20 million cells were enough. For TF ChIP 20-30 million cells were used. For transcription factor ChIP, 30 million epidermal cells were cross-linked with 1/250 of Gold fixative (C01019027, Diagenode) in 30 ml of PBS with 1mM of MgCl_2_, with incubation for 30 min with mild swirling at room temperature.

After two PBS washes, a second fixation was performed in methanol-free 1% formaldehyde (15710, Electron Microscopy Sciences) in 30 ml of MEM calcium-free medium (M8167, Sigma) with 10% calcium-chelated fetal bovine serum (FBS, A5256801, Life Technologies; Chele× 100 Chelating Resin, 1421253, Biorad) for 10 min under rotation at room temperature. Straight after, glycine (50046, Sigma) was added to a final concentration of 125 mM to stop cross-linking with incubation for 5 min under rotation at room temperature. For chromatin mark ChIP, 20 million epidermal cells were cross-linked in a single fixation step with methanol-free 1% formaldehyde in MEM calcium-free medium with 10% calcium-chelated fetal bovine serum, following the same steps explained above for transcription factor ChIP. From here onwards, the same steps were followed for both histone mark and transcription factor ChIPs. The pellet was washed twice with cold PBS and resuspended after with 7.5 ml of swelling buffer (25mM HEPES pH 7.9, 1.5mM MgCl_2_, 10 mM KCl, 0.1% NP-40 supplemented with 1X protease inhibitors without EDTA), and incubated for 10 min on ice. This cell suspension was homogenised with a Dounce homogenizer and tight pestle (885302-0015, Kimble) 50 times. Cell extracts were then centrifuged for 5 min at 3000g at 4°C. Pelleted nuclear extracts were resuspended (in 1/10 of the volume used for the swelling buffer) in ChIP buffer (10 mM Tris-HCl pH 7.5, 150mM NaCl, 1% Triton X-100, 5mM EDTA pH 8.0, 0.5 mM DTT, supplemented with 0.2% SDS - in the case of Gold fixated chromatin, supplement with 0.5% SDS for non-Gold fixated chromatin - and 1X protease inhibitors without EDTA), and incubated for 15 min on ice. This suspension was then transferred to sonication tubes (C01020031, Diagenode) and sonication was carried out in 30 cycles of 30/30 min on/off at 4°C (Bioruptor Pico, Diagenode). Sonicated material was centrifuged at 14000 g for 10 min at 4°C. The supernatant contains the chromatin, which was quantified for ChIP-seq.

For immunoprecipitation, 30 ug of chromatin were used for transcription factor ChIP while 20 ug were used for chromatin mark ChIP. As the input, a volume corresponding to 1% of the amount of chromatin used for ChIP was taken and transferred to another tube to keep at -20°C until used. Chromatin for immunoprecipitation was diluted with ChIP buffer without SDS to dilute the SDS concentration to 0.1% in all samples. The following antibodies and final concentrations were used for the pulldown of the different transcription factors and chromatin marks: 10 ug ml^−1^ anti-BMAL1 (ab3806, Abcam), 20 ul anti-YAP (CS14074, Cell Signaling Technology) for 20 ug of chromatin, 5 ul anti-p65 (8242, Cell Signaling Technology) for 10 ug of chromatin, 2 ul anti-H3K27ac (ab4729, Abcam) for 25 ug of chromatin, 2 ul anti-H3K4me1 (ab8895, Abcam) for 25 ug of chromatin, 3.25 ug anti-H3K4me3 (C15410003, Diagenode) for 20 ug of chromatin.

Samples were incubated with rotation with the corresponding antibody overnight at 4°C. Protein A Sepharose beads (17-5280-01, Merck) were used according to the manufacturer’s instructions, added to the sample tubes and incubated with rotation for 2 hours at 4°C. After, samples were centrifuged at 1000g for 3 min at 4°C and washed with 1 ml of low-salt (50mM HEPES pH 7.5, 140 mM NaCl, 1% Triton X-100, supplemented with 1X protease inhibitors without EDTA) and high-salt (50mM HEPES pH 7.5, 500 mM NaCl, 1% Triton X-100, supplemented with 1X protease inhibitors without EDTA) buffers. Then, a final wash with TE buffer (10 mM Tris-HCl pH 8.0, 1mM EDTA pH 8.0) was performed. Immunoprecipitated chromatin was eluted in elution buffer (1% SDS, 100mN NaHCO_3_, freshly prepared) by incubation with the elution buffer for 30 min at 65°C with agitation. Then, samples were centrifuged at 1000g for 3 min, and the supernatant containing the DNA was transferred to a new tube. 5M NaCl was added to a final concentration of 200mM to the eluted DNA and input DNA, followed by incubation at 65°C overnight with agitation. On the following day, Tris-HCl pH 6.8 was added to a final concentration of 40mM, EDTA pH 8.0 to a final concentration of 10mM and proteinase K (3115879001, Merck) to a final concentration of 50 ug ml^−1^, followed by incubation for 1 hour at 45°C. Lastly, DNA was purified using the QIAquick PCR Purification Kit (28106, Qiagen).

### ChIP-seq library preparation and sequencing

Libraries were carried out using the NEBNext Ultra DNA Library Prep for Illumina kit (E7370) or the NEBNext® Ultra II DNA Library Prep for Illumina® kit (E7645) using the manufacturer’s protocol. In brief, ChIP-enriched DNA and input DNA were subjected to end repair and the addition of “A” bases to 3’ ends, NEB adapter’s ligation and USER excision. For all purification steps, AgenCourt AMPure XP beads (A63882, Beckman Coulter) were used. Library amplification was performed by PCR using NEBNext Multiplex Oligos for Illumina (96 Unique Dual Index Primer Pairs, E6440, E6442, E6444, E6446). Analysis of final libraries was carried out using either an Agilent Bioanalyzer or Fragment analyzer High Sensitivity assay (5067-4626 or DNF-474) to estimate the quantity and check size distribution. Quantification of these final libraries was done by quantitative PCR using the KAPA Library Quantification Kit (KK4835, KapaBiosystems, 07960204001 Roche). Libraries were sequenced 1 x 51 + 10 + 10 base pairs on Illumina’s NextSeq2000. Between 25 and 50 million reads were obtained per sample.

### ChIP-seq data processing and Deeptools clustering

Reads were trimmed, adapters were removed and low-quality reads were discarded by using Trimmomatic^103^ (v.0.36, TRAILING:5 SLIDINGWINDOW:4:15 MINLEN:36). Reads were then aligned to the mm10 genome with Burrows-Wheeler aligner^104^ (v.0.7.12, -n 2 -l 20 -k 1 -t 2). Until this point, each sample was processed individually. Then, aligned reads (.bam files) from the biological replicates were merged and duplicates were removed using SAMtools^105^ (v.1.5). Downsampling to 100 million reads was performed for BMAL1 and to 60 million reads for YAP and chromatin marks to obtain a pool of three or four replicates per condition. The same downsampling was done for the corresponding input DNAs. Peaks were called in these pooled files using MACS2^106^ (v.2.1.1) with the following parameters: -q 0.01 -nomodel -extsize 300 -B -SPMR, except for H3K4me1 where broad peaks were called with the -broad option. Only peaks with a peak score >300 were considered for YAP ChIP analysis; peaks with >200 were chosen for BMAL1, and peaks with >20 for the rest of ChIPs.

H3K27ac, H3K4me and H3K4me3 ChIP-seq signal profiles were measured in the BMAL1 or YAP peak regions, followed by unsupervised clustering using Deeptools^107^. Heat maps of these ChIP-seq signal profiles clustering were generated using Deeptools 3.2.1 functions bamCoverage (options -bs 50 --smoothLength 150 --normalizeUsing CPM), bigwigCompare, computeMatrix and plotHeatmap.

Peaks were annotated to the closest gene with Homer version 4.11^108^. Only peaks annotated to protein-coding genes were used for Gene Ontology (GO) analysis.

### Permutation test to assess ChIP-seq signal overlap

To evaluate the statistical significance of the association between BMAL1 and YAP peaks, a permutation test was performed with RegioneR using 5000 permutations^83^.

### Motif enrichment analysis

Transcription factor motif discovery analysis was performed with HOMER^108^ (v.4.11) function *findmotifs*. Default parameters were used. The results of selected motifs found in the *de novo* enrichments are presented.

### Gene Ontology enrichment analysis

Gene Ontology (GO) analysis for RNA-seq and ChIP-seq data was performed by using the Enrichr web tool^109^ (https://maayanlab.cloud/Enrichr/). A category was considered significant if *P*-value < 0.01.

### Immunofluorescence (IF) on FFPE skin sections

For immunofluorescence from murine skin, samples of back skin were taken, fixed in neutral buffered formalin (10%) for 3 h at room temperature, dehydrated and embedded in paraffin blocks. Formalin-fixed paraffin-embedded (FFPE) blocks were cut into 2-4 µm sections.

Depending on the primary antibody used the immunofluorescence protocol conditions vary, as indicated in the following paragraph.

Anti-YAP1 phospho Y357: following heat-mediated antigen retrieval (20 min at 97°C with citrate pH 6.0), sections were blocked with 0.25% gelatine in PBS for 90 min at room temperature. Incubation of the primary antibody anti-YAP1 (phospho Y357) (ab62751, Abcam) in 1:400 in Dako Antibody Diluent with Background Reducing Components (Dako, S3022) was performed overnight at 4°C. After three washes with 0.2% Tween20 0.25% gelatine in PBS, incubation of the secondary antibody Alexa Fluor^TM^ 488 donkey anti-rabbit IgG (H+L) (A21206, Invitrogen) 1:400 in Dako Antibody Diluent with Background Reducing Components (Dako, S3022) was performed at room temperature for 1 h. After three washes with 0.2% Tween20 0.25% gelatine in PBS, nuclei were counterstained with 0.5 ug ml^−1^ of DAPI for 10 min at room temperature. Slides were mounted with Dako Fluorescence Mounting Medium (Dako, S3023). 5 aged and 5 adult mice were analysed for the aged vs adult comparison. 5 aged and 5 adult mice were analysed for the aged vs adult comparison. 5 anti-IL-17A/F treated aged mice and 7 IgG control-treated aged mice were analysed for the anti-IL-17A/F comparison. 8 ColXIV KO and 7 ColXIV WT 47-week-old mice were analysed.

Anti-unphosphorylated Ser127 YAP1 (unP-S127): following heat-mediated antigen retrieval (20 min at 97°C with Tris EDTA pH 9.0), sections were blocked with 10% donkey serum in PBS for 1 h at room temperature. Incubation of the primary antibody (anti-active YAP1, ab205270, Abcam, 1:1000 in Dako Antibody Diluent with Background Reducing Components, Dako, S3022) was performed overnight at 4°C. After three washes with PBS, incubation of the secondary antibody (Alexa Fluor^TM^ 488 donkey anti-rabbit IgG (H+L), A21206, Invitrogen, 1:400 in Dako Antibody Diluent with Background Reducing Components, Dako, S3022) was performed at room temperature for 1 h. After three washes with PBS, nuclei were counterstained with 0.5 ug ml^−1^ of DAPI for 10 min at room temperature. Slides were mounted with Dako Fluorescence Mounting Medium (Dako, S3023). 12 aged and 10 adult mice were analysed for the aged vs adult comparison. 8 ColXIV KO and 7 ColXIV WT 47-week-old mice were analysed. 7 anti-IL-17A/F treated aged mice and 8 IgG control-treated aged mice were analysed for the anti-IL-17A/F comparison.

### Proximity Ligation Assay (PLA)

Formalin-fixed paraffin-embedded (FFPE) blocks were cut into 2-4 µm sections. Deparaffinization and heat-mediated antigen retrieval for 20 min at 97°C with Citrate pH 6.0 was performed before PLA reaction protocol. For PLA reaction, NaveniFlex Tissue MR Atto647N (NT.MR.100 Atto647N, Navinci) was used following the manufacturer’s instructions. Incubation of the primary antibodies (anti-BMAL1 (ab3350, Abcam) at 1:200 and anti-YAP (12395, Cell Signalling) at 1:100, diluted in the diluent provided by PLA manufacturer’s kit) was performed at °4C overnight. A test with only the Rabbit antibody was performed to exclude the possibility of endogenous mouse IgG performing unspecific PLA signal with the rabbit secondary antibody.

### Image acquisition

Full images were acquired with either a NanoZoomer-2.0 HT C9600 digital scanner (Hamamatsu) with the 20X objective, in which 1 pixel corresponds to 0.46um, and coupled to a mercury lamp unit L11600-05 and using NDP.scan2.5 software U10074-03 (Hamamatsu, Photonics, France) or Phenoimager HT digital scanner (Akoya Biosciences) with a 20X objective. For the representation of PLA, images were captured with Zeiss LSM 780 confocal with an immersion 63x objective. Brightness and contrast settings were enhanced for visualization purposes exclusively.

### Histopathological analysis

Scaled images were analysed with Qupath v0.3.0 (for IF) or v0.5.0 (for PLA samples). For immunostainings, multiple selections (region of interest - ROI) of the mouse skin IFE, excluding folded areas and the HFs, were done manually in two portions of the skin. The “Positive cell count” algorithm was used to detect positive cells in the 488-channel. For PLA, nuclei were detected and the subcellular spot detection function was used to count speckles per nucleus on the Cy5 channel.

### Atomic force microscopy

Atomic force microscopy measurements of interfollicular BMs were performed on freshly cut 16-micron cryosections using JPK Nanowizard 2 system (Bruker Nano) mounted onto an Olympus IX73 microscope (Olympus). Briefly, cryosections were equilibrated in 1X PBS supplemented with 1X protease inhibitors and measured within 20 minutes post-thaw. Triangular nonconductive silicone nitride MLCT cantilevers (Bruker) with a nominal spring constant of 0.01Nm-1 were used for nanoindentation. Forces of 2nN were applied. Analyses were carried out using JPK SPM v5 software by fitting the Hertz model corrected for tip geometry post removal of offset from baseline, identification of contact point, and subtraction of cantilever bending to obtain Young’s Modulus of samples.

### *In vivo* anti-IL-17A/F neutralising treatment

Cohorts of aged (80 weeks old) mice randomly distributed into two groups and were treated with either a mixture of 105 ug of anti-IL-17A (clone 17F3, BE0173, BioXCell) and 105 ug of anti-IL-17F (clone MM17F8F5.1A9, BE0303, BioXCell) for anti-IL-17A/F treatment group, or 210 ug of control IgG1 (clone M0PC-21, BE0083, BioXCell) for the control group. Injections of 100 ul of antibody solution in PBS were administered intraperitoneally and performed thrice weekly. Mice were treated for 4 weeks, and then mice were sacrificed at ZT4 to obtain samples for genomic and histologic analysis.

### Statistical analysis

Generally, graphs show individual values and median (depicted as a line). *P-*values were obtained with the Mann-Whitney *U* test with Prism v9 unless otherwise stated in the figure legend.

### Data availability

Sequencing data is available at GSE294389 (https://www.ncbi.nlm.nih.gov/geo/query/acc.cgi?acc=GSE294389 ; token ehcrqmakpjenheh).

## Acknowledgements

Research in the S.A.B. lab is supported partially by the European Research Council (ERC) under the European Union’s Horizon 2020 research and innovation programme (Grant Agreement No. 787041), the Government of Cataluña (SGR grant), the Government of Spain (MINECO), the La Marató/TV3 Foundation, the Foundation Lilliane Bettencourt, the Spanish Association for Cancer Research (AECC) and The Worldwide Cancer Research Foundation (WCRF). P.S. was awarded a Spanish Ministry of Economy and Development fellowship (MINECO; ID BES-2017-081279).

J.B. was awarded a Spanish Ministry of Science and Innovation fellowship (MCI; ID PRE-2021-097118). The IRB Barcelona is a Severo Ochoa Centre of Excellence (MINECO award SEV-2015-0505). We would like to thank the Histopathology, Advanced Digital Microscopy and Functional Genomics units at the IRB Barcelona for their assistance in this work. We would like to thank the genomics unit at the CRG for assistance with sequencing.

## Author contributions

P.S., J.B. and G.S. designed the experiments. P.S., J.B., G.S. and L.A. performed the experiments and collected data. P.S., J.B. and G.S. analysed and interpreted the data. G.S., P.S., J.B., O.R., T.M. and C.S.A. carried out bioinformatics analyses. Y.A.M. and S.A.W. performed atomic force microscopy measurements. J.B., S.A.B. and G.S. drafted the manuscript. G.S. and S.A.B. conceived the study and supervised the project. All authors contributed to the final version of the manuscript.

---

**Extended Data Fig.1.**
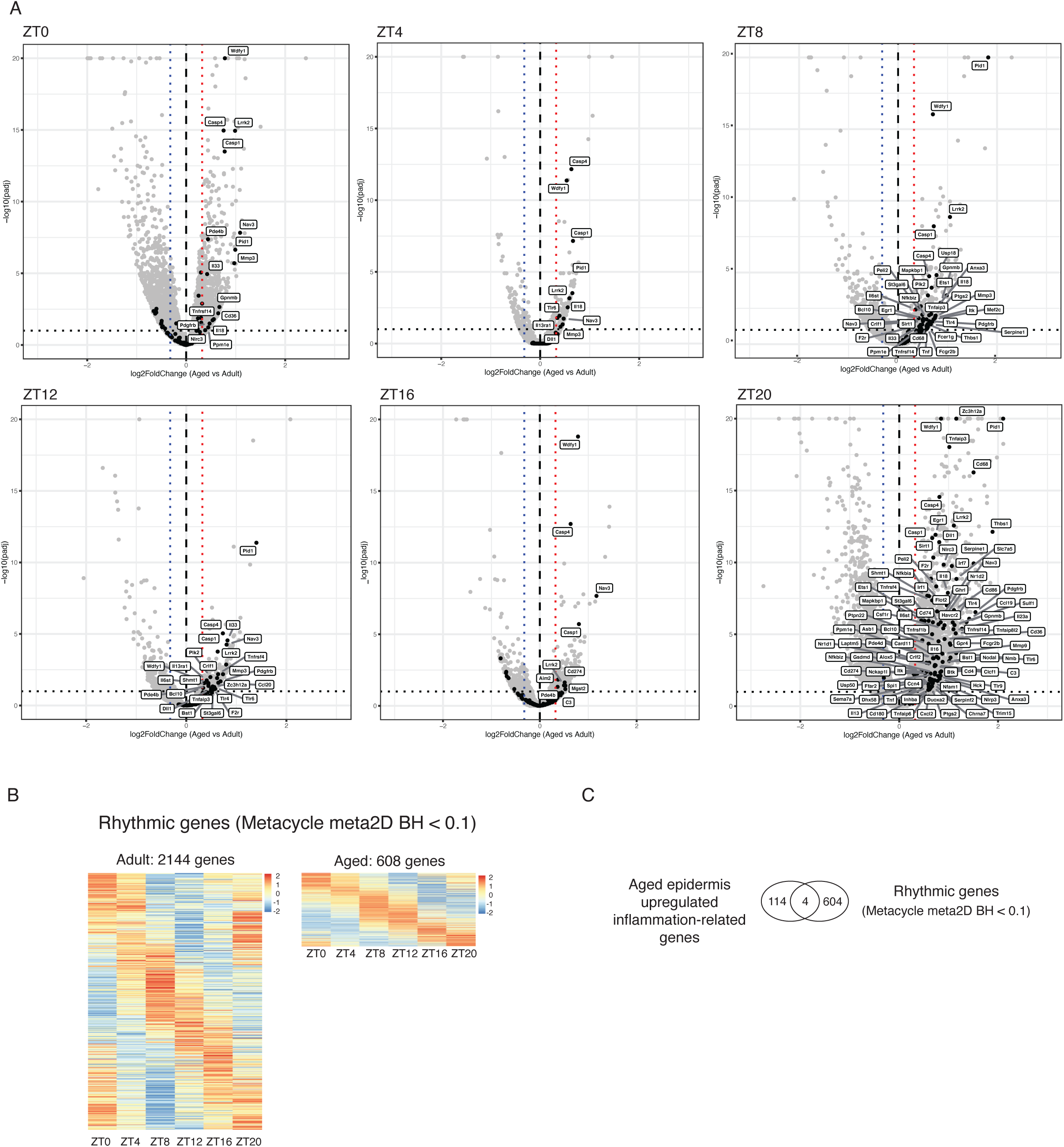
Up-regulation of inflammation-related genes during ageing does not happen rhythmically. **A.** DEGs between adult and aged epidermis at each ZT. Inflammation-related genes that are up-regulated in at least one ZT are highlighted in black. **B.** Heatmap representation of rhythmic gene expression throughout the ZTs in adult and aged epidermis obtained with Meta2D analysis. Only genes with a BH< 0.1 have been considered statistically significant rhythmic genes. **C.** Comparison of the inflammation-related genes up-regulated during ageing with rhythmic genes during ageing. ZT, Zeitgeber time.

**Extended Data Fig.2.**
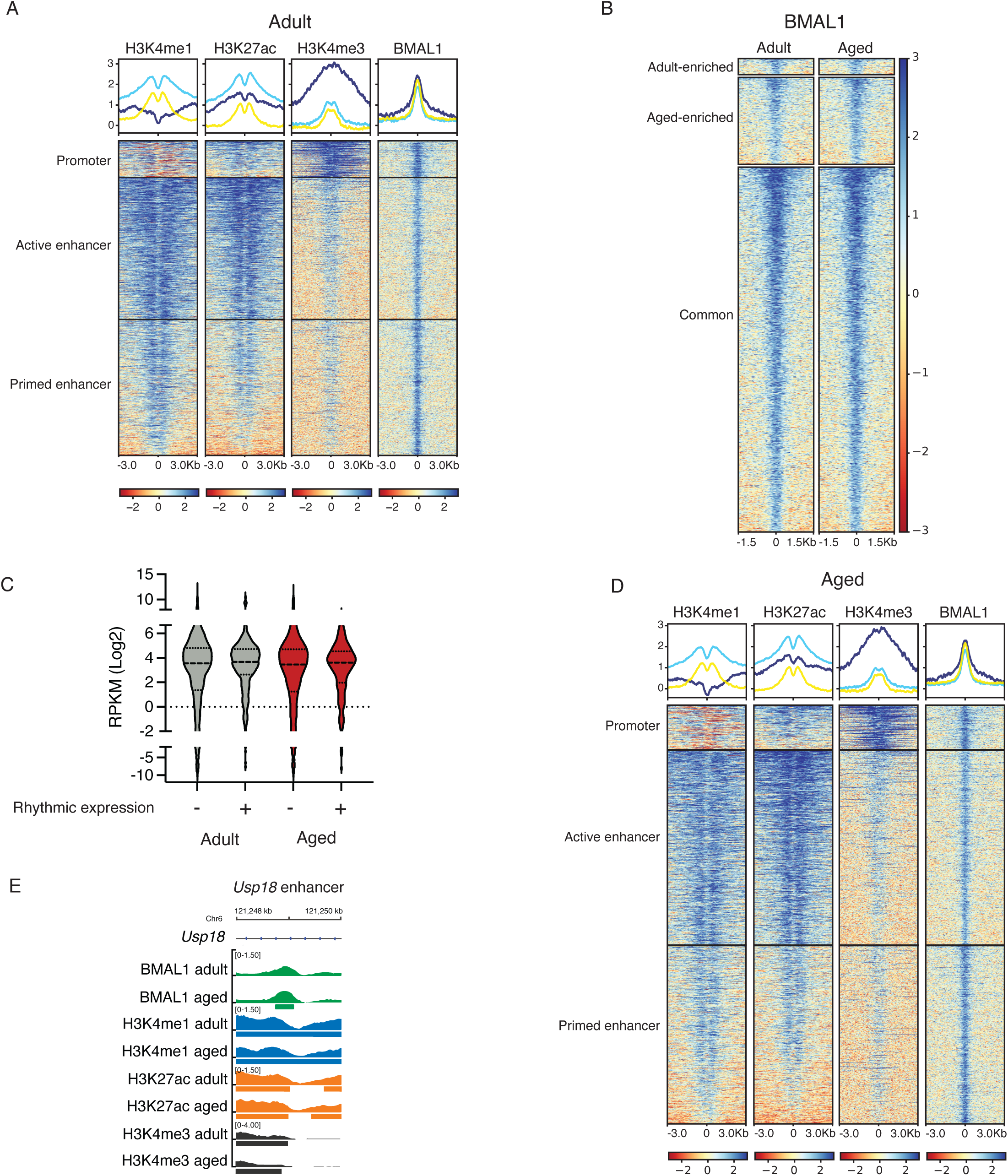
BMAL1 genomic occupancy in adult and aged epidermis. **A.** Heatmap and density plots of H3K4me1, H3K27ac, H3K4me3 and BMAL1 ChIP-seq signal score ± 3 kb of peak centre at BMAL1 individual peaks in adult epidermis. K-means (k=3) clustering of BMAL1 peaks was performed based on H3K4me1, H3K27ac, H3K4me3 and BMAL1 signal score. **B.** Heatmap of BMAL1 ChIP-seq signal score ± 1.5 kb of peak centre of individual peaks in adult and aged epidermis. K-means (k=3) clustering of BMAL1 peaks was performed based on BMAL1 signal score. **C**. Violin plot showing expression values (RPKM) of rhythmic and non-rhythmic BMAL1 targets in adult and aged epidermis. **D.** Heatmap and density plots of H3K4me1, H3K27ac, H3K4me3 and BMAL1 ChIP-seq signal score ± 3 kb of peak centre at BMAL1 individual peaks in aged epidermis. K-means (k=3) clustering of BMAL1 peaks was performed based on H3K4me1, H3K27ac, H3K4me3 and BMAL1 signal score. **E.** Genomic loci depicting BMAL1 binding to an enhancer region of *Usp18*, one of the inflammation-related genes whose expression is up-regulated in aged epidermis. The thick bars under genomic tracks represent those that have been called a peak for the TF or chromatin mark. Chromosome kilobases are displayed above for identification of the loci. RPKM, Reads Per Kilobase per Million mapped reads.

**Extended Data Fig.3.**
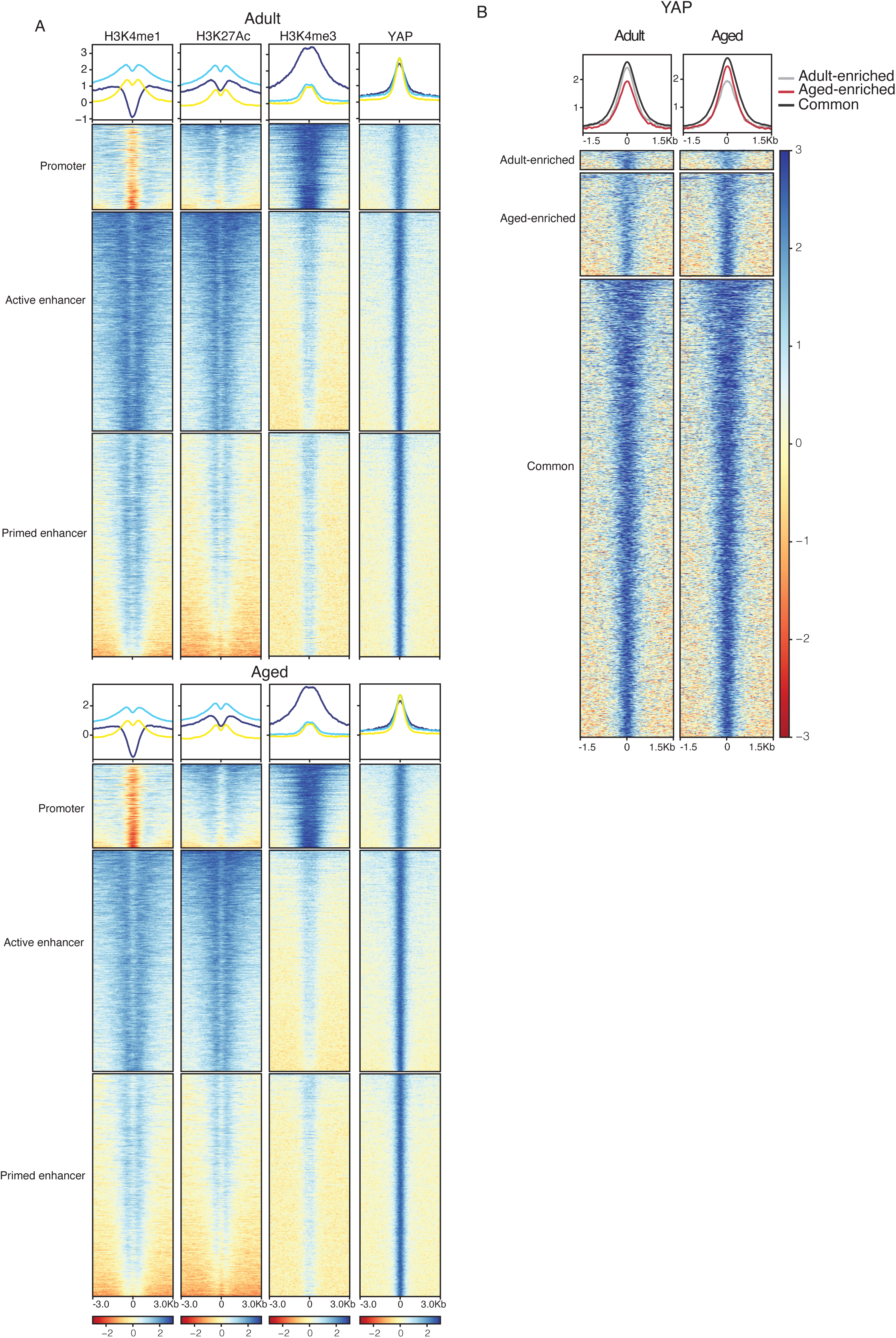
YAP genomic occupancy in adult and aged epidermis. **A.** Heatmap and density plots of H3K4me1, H3K27ac, H3K4me3 and YAP ChIP-seq signal score ± 3 kb of peak centre at YAP individual peaks in adult (upper) and aged (lower) epidermis. K-means (k=3) clustering of YAP peaks was performed based on H3K4me1, H3K27ac, H3K4me3 and YAP signal scores. **B.** Heatmap and density plots of YAP ChIP-seq signal score ± 1.5 kb of peak centre of individual peaks in adult and aged epidermis. K-means (k=3) clustering of YAP peaks was performed based on YAP signal score.

**Extended Data Fig.4.**
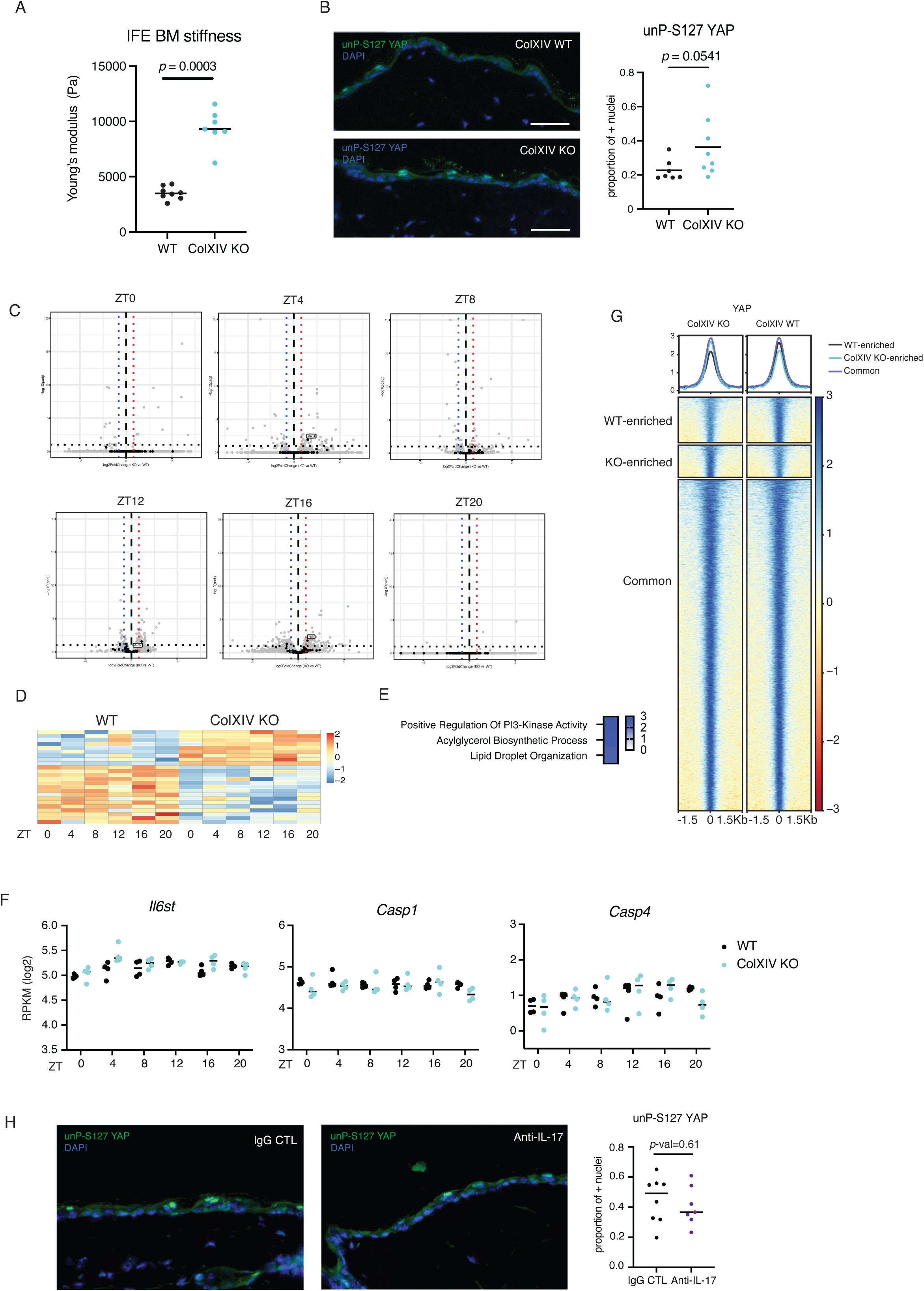
Increased IFE BM stiffness in the ColXIV KO mouse model does not recapitulate aged-related gene expression nor YAP genomic occupancy. **A.** Atomic force microscopy force indentation measurements of IFE BM stiffness. Between 20-100 force curves were taken and the mean (represented as a dot) was calculated for each mouse. n = 7 ColXIV KO and n = 8 ColXIV WT mice at 47 weeks old. **B.** Representative images of immunofluorescence staining of unphosphorylated-S127 YAP (unP-S127 YAP) and quantification in ColXIV WT and ColXIV KO mouse back skin. Scale bars, 50 μm. For enhanced visualization, brightness and contrast were adjusted on these images for visualization purposes exclusively, and equally in both conditions; n= 7 ColXIV WT mice and n= 8 ColXIV KO mice at 47 weeks old. All quantifications are presented as the proportion of epidermal cell nuclei positive for staining. **C.** DEGs between ColXIV WT and ColXIV KO epidermis at each ZT. Inflammation-related genes that are up-regulated during ageing (from the aged vs adult epidermis comparison) in at least one ZT are highlighted in black. **D.** Heatmap showing expression levels of DEGs in ColXIV WT and ColXIV KO epidermis throughout the day (6 ZTs). Transcriptomes of epidermal cells from n= 4 ColXIV WT and n= 4 ColXIV KO mice. **E.** GO BP enrichment analysis of up-regulated genes in ColXIV KO epidermis compared to WT irrespective of their rhythmicity pattern. Only selected GO BP categories are shown. The colour intensity represents the -log_10_ of the *p*-value for each depicted category. **F.** *Il6st*, *Casp1* and *Casp4* gene expression throughout the day in ColXIV WT and ColXIV KO epidermis. The y-axis represents the RPKM for each replicate and x-axis shows ZTs **G.** Density plots and heatmap of YAP ChIP-seq signal ± 1.5 kb of peak centre in ColXIV KO and ColXIV WT epidermis at 47 weeks old. K-means (k=3) clustering of YAP peaks was performed based on YAP signal score. Colours depict the signal of the TF in each cluster (black= ColXIV WT enriched, light blue= ColXIV KO enriched and blue=common) of YAP peaks. **H.** Representative images of immunofluorescence staining of unphosphorylated-S127 YAP (unP-S127 YAP) and quantification in anti-IL-17 and IgG control aged mouse back skin. Scale bars, 50 μm. n= 7 anti-IL-17 and n= 8 control IgG mice. IFE, interfollicular epidermis; BM, basement membrane; GO BP, Gene Ontology Biological Processes; ZT, Zeitgeber time; RPKM, Reads Per Kilobase per Million mapped reads.

**Extended Data Fig.5.**
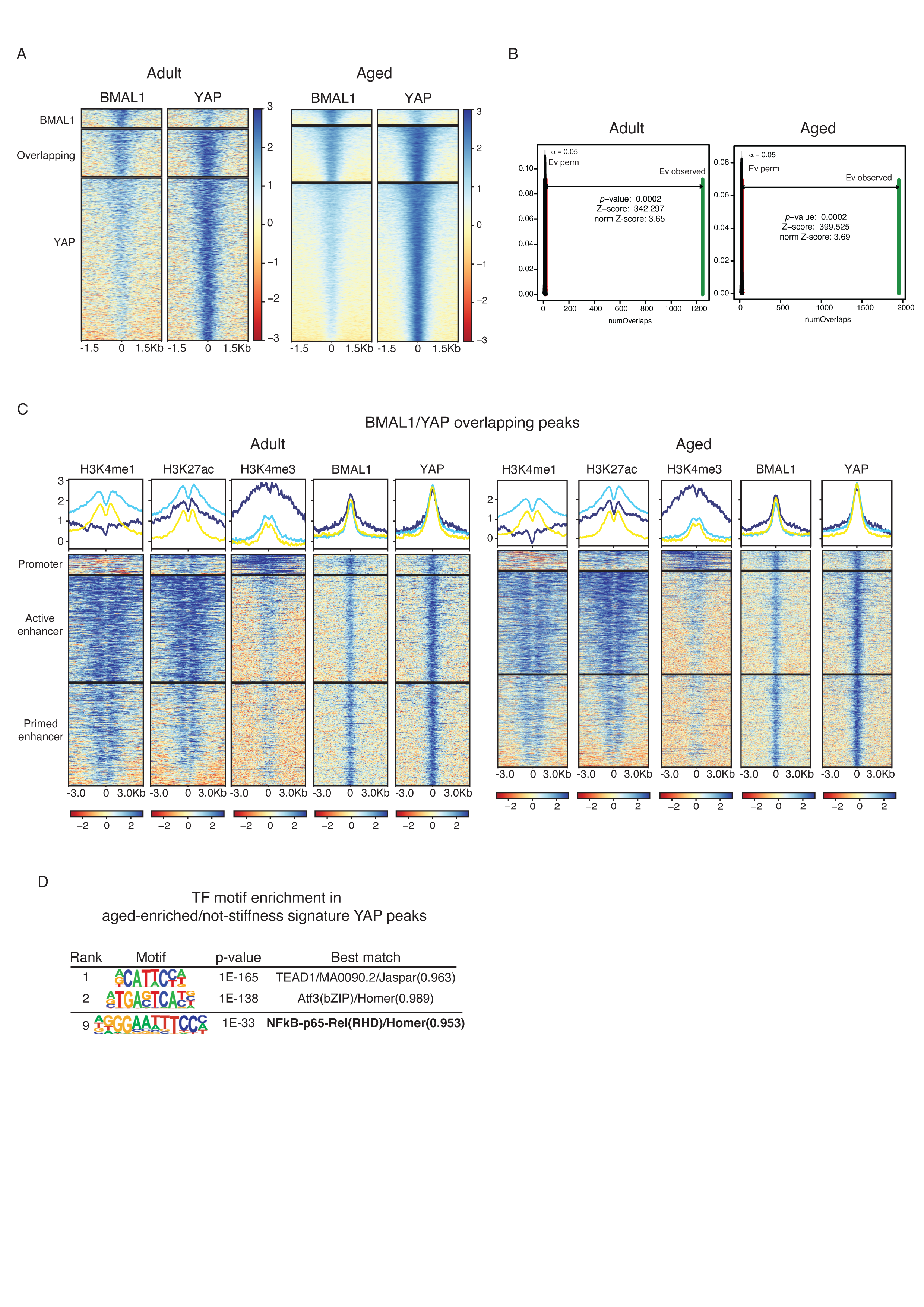
BMAL1 and YAP share genomic binding to a set of CREs from adult and aged epidermis. **A.** Heatmap of BMAL1 and YAP ChIP-seq signal score ± 1.5 kb of peak centre of individual peaks in adult (left) and aged (right) epidermis. K-means (k=3) clustering of BMAL and YAP peaks was performed based on their signal score. **B.** Permutation test results evaluating the association between BMAL1 and YAP peaks. The BMAL1–YAP peaks association is highly significant, with an observed overlap value far from the value obtained by random distribution. Green lines indicate the observed overlap of BMAL1 and YAP peaks, and grey lines indicate the significance threshold in an expected random distribution. **C.** Heatmap and density plots of H3K4me1, H3K27ac, H3K4me3, BMAL1 and YAP ChIP-seq signals ± 3 kb of peak centre at BMAL1 and YAP overlapping peaks in adult (left) and aged (right) epidermis. K-means (k=3) clustering of BMAL1 and YAP overlapping peaks was performed based on H3K4me1, H3K27ac, H3K4me3, BMAL1 and YAP signal score. **D.** TF motifs enriched in aged-enriched/not stiffness signature YAP peaks, based on de *novo* motif discovery performed with HOMER. TF, transcription factor.

**Extended Data Fig.6.**
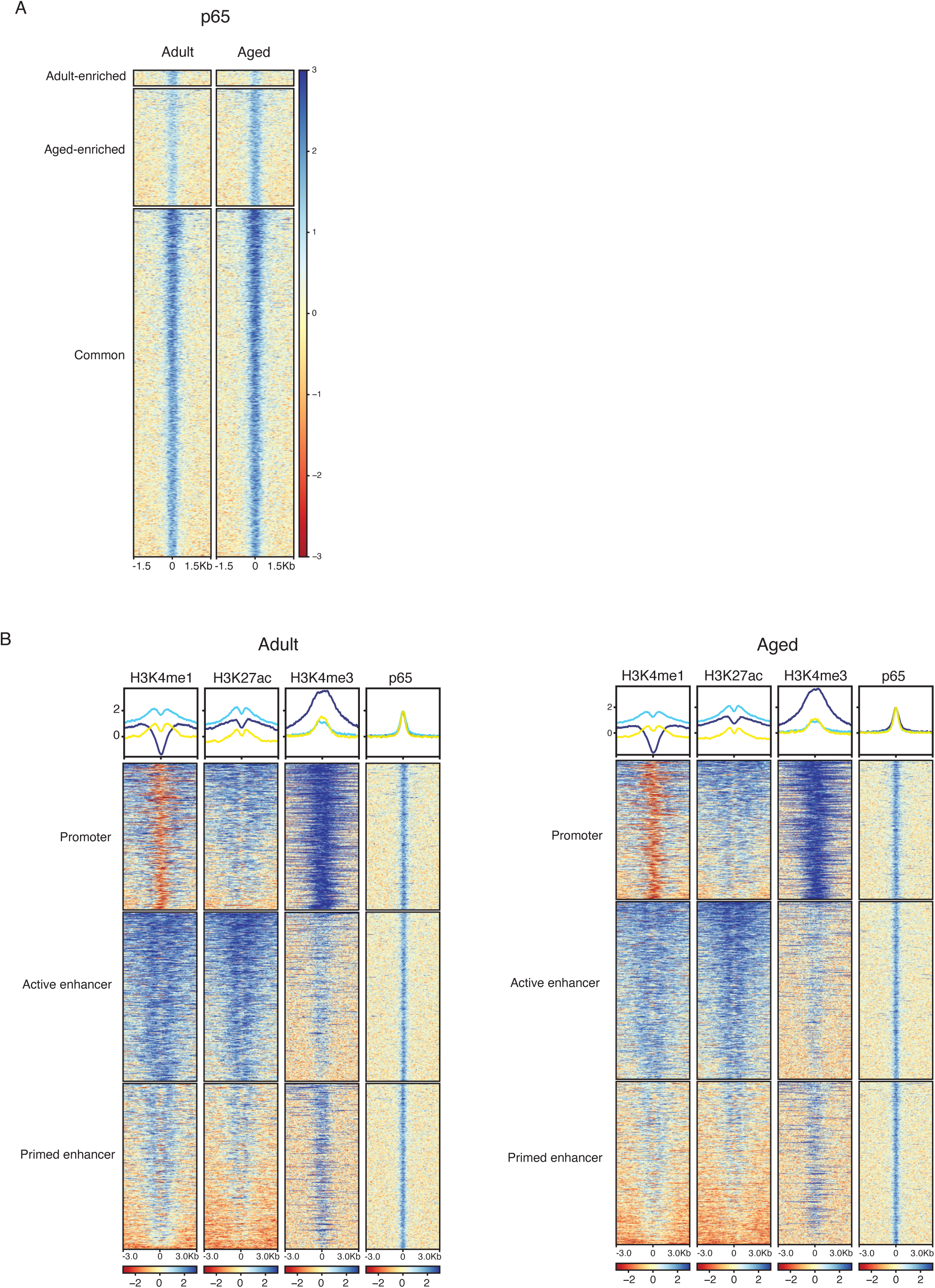
p65 genomic occupancy in adult and aged epidermis. **A.** Heatmap of p65 ChIP-seq signal score ± 1.5 kb of peak centre of individual peaks in adult and aged epidermis. K-means (k=3) clustering of p65 peaks was performed based on p65 signal score. **B.** Heatmap and density plot of H3K4me1, H3K27ac, H3K4me3 and p65 ChIP-seq signal score ± 3 kb of peak centre at p65 individual peaks in adult (left) and aged (right) epidermis. K-means (k=3) clustering of p65 peaks was performed based on H3K4me1, H3K27ac, H3K4me3 and p65 signal scores.

